# Revealing molecular determinants of ligand efficacy and affinity at the D_2_ dopamine receptor through molecular dynamics simulations

**DOI:** 10.1101/2025.03.30.646192

**Authors:** Yue Chen, Nour Aldin Kahlous, Marcus Saarinen, Sergio Perez Conesa, Per Svenningsson, Lucie Delemotte, Jens Carlsson

## Abstract

G protein-coupled receptors (GPCRs) control numerous physiological processes and are important therapeutic targets. Despite major research efforts, rational design of drugs that stimulate GPCR signaling is challenging because the molecular basis of activation remains poorly understood. Here, a combination of molecular dynamics simulations and pharmacological assays was used to study the activation mechanism of the D_2_ dopamine receptor (D_2_R), a major drug target for central nervous system diseases. Enhanced sampling simulations were performed to identify the key conformational changes involved in D_2_R activation by dopamine, and a computational platform for ligand profiling based on free energy calculations was developed. Simulations and experimental characterization of a series of dopamine derivatives showed that free energy calculations can predict the effect of small chemical modifications on ligand affinity and efficacy. Furthermore, simulations of D_2_ dopamine and β_2_ adrenergic receptor activation revealed that ligand-induced activation of these GPCRs is driven by different molecular mechanisms despite recognizing chemically similar catecholamine agonists. Whereas dopamine interactions with the sixth transmembrane helix primarily drive activation of the D_2_R, hydrogen bonding with the fifth helix is the key interaction for activation of the β_2_ adrenergic receptor. Our results highlight the complexity of GPCR activation and illustrate how molecular simulations can provide mechanistic insight and quantitative predictions of ligand activity, enabling structure-based drug design.

**Significance statement:** G protein-coupled receptors (GPCRs) on the cell surface recognize ligands such as hormones and neurotransmitters. The binding of the ligand to the receptor leads to the activation of intracellular signaling pathways, a communication system that controls essential physiological functions. GPCRs are therefore important therapeutic targets, and many drugs exert effects by modulating their activity. Here, we use molecular simulations to study the activation mechanism of a dopamine receptor, which is implicated in neurodegenerative and neuropsychiatric disorders. We elucidate the mechanism of receptor activation and develop strategies to predict ligand affinity and efficacy. We also identify differsences in the activation mechanisms of dopamine and adrenergic receptors. These findings provide novel insights into receptor activation and can accelerate the drug discovery process.

## Introduction

The family of G protein-coupled receptors (GPCRs) regulates cellular responses to signaling molecules such as neurotransmitters and hormones by activating intracellular effectors, including heterotrimeric G proteins and arrestins. Because of the essential roles of GPCRs in physiology, members of this large superfamily are implicated in a wide range of diseases and have become important drug targets (1, 2). Development of ligands that control GPCR activity by either stimulating or blocking signaling has contributed to the understanding of physiological processes and led to the discovery of numerous drugs. However, rational drug design remains challenging because the molecular mechanism of receptor activation is poorly understood.

Crystal and cryo-EM structures of GPCRs in complex with ligands and effectors have revealed drug binding sites and conformational changes involved in activation (3, 4). In class A receptors, the extracellular binding site is located in the middle of the transmembrane region, which contains the seven helices (TM1-7) characteristic of GPCRs. In the intracellular region, the major conformational change is an outward movement of TM6, which enables binding of G proteins or arrestins. NMR spectroscopy has revealed that GPCRs are flexible proteins and occupy both active- and inactive-like conformations in the absence of bound ligand, resulting in basal activity (3, 5). Ligand binding to the orthosteric site can perturb this equilibrium, leading to a spectrum of efficacy profiles. Agonists favor the ensemble of active conformations whereas inverse agonists stabilize inactive states. Antagonists do not influence the equilibrium between active and inactive states, and partial agonists activate the receptor but do not reach the maximal efficacy. Intriguingly, ligands with different effects on receptor signaling have similar chemical structures and binding modes, and small modifications can lead to large changes in pharmacological profile (e.g., affinity and efficacy) (6, 7). These effects illustrate the complexity of GPCR activation and are difficult to explain solely based on the static experimental structures. Compounds with improved potency and tailored efficacy profiles could contribute to the development of more efficient drugs with less side effects. However, design of such drugs will require a better understanding of how ligand binding influences the conformational landscape of the receptor.

The aminergic receptors are the most extensively characterized group of GPCRs and are activated by biogenic amines such as adrenaline, dopamine, serotonin, histamine, or trace amines. These receptors play essential roles in numerous physiological processes and constitute the vast majority (∼70%) of the established GPCR drug targets (2). Representative atomic resolution structures demonstrating how the biogenic amines bind to their cognate receptor are now available and have revealed that their orthosteric sites and receptor-ligand interactions are highly similar (8). For example, the five dopamine and nine adrenergic receptors in humans recognize closely related catecholamines and most of the ligand-contacting residues are conserved. These observations highlight why most drugs targeting aminergic GPCRs also have side effects caused by undesired interactions with other subtypes (9). More detailed insight into the activation mechanisms of adrenergic and dopamine receptors could facilitate the design of subtype selective ligands.

In this work, we combined structure-based modeling and pharmacological assays to elucidate the activation mechanism of the D_2_ dopamine receptor (D_2_R), a major drug target for central nervous system diseases such as Parkinson’s disease and schizophrenia (10, 11). Molecular dynamics (MD) simulations have the potential to provide detailed information on the allosteric modulation of GPCRs by ligands at the molecular level, but the time scales of these processes are longer than what can be achieved with the available computational resources. Enhanced sampling techniques and alchemical free energy calculations have been shown to be effective approaches to study receptor activation and ligand binding (12–17). Here, we applied these techniques to elucidate the mechanism of D_2_R activation and develop a computational platform for efficient profiling of the efficacy and affinity of ligands. Based on pharmacological assays of a series of compounds closely related to dopamine, predictive models based on MD simulations were obtained and revealed unique aspects of D_2_R activation that can be utilized in drug discovery.

## Results

### Molecular mechanism of D_2_R activation by dopamine

Several structures of the D_2_R in inactive and active conformations were available, which enabled identification of residues involved in the G protein activation (18–23). The transition from inactive to active states involves the reorganization of several microswitches (molecular motifs conserved in GPCRs) in the transmembrane (TM) region (Figure 1a). In the orthosteric site, receptor activation involves conformational changes in the CWxP motif in TM6 (C385^6.47^, W386^6.48^, and P388^6.50^; superscripts represent Ballesteros-Weinstein numbering (24)). This rearrangement is accompanied by the formation of an active PIF motif (P201^5.50^, I122^3.40^, and F382^6.44^) below the binding site (the connector region) and an inward twist of the NPxxY motif in TM7 (N422^7.49^, P423^7.50^, and Y426^7.53^), leading to an interaction between Y426^7.53^ and Y209^5.58^ (YY motif). These conformational changes disrupt the ionic lock (a salt bridge between E368^6.30^ and R132^3.50^ in the DRY motif), and an outward movement of TM6 in the intercellular region leads to formation of the G protein binding site.

**Figure 1.**
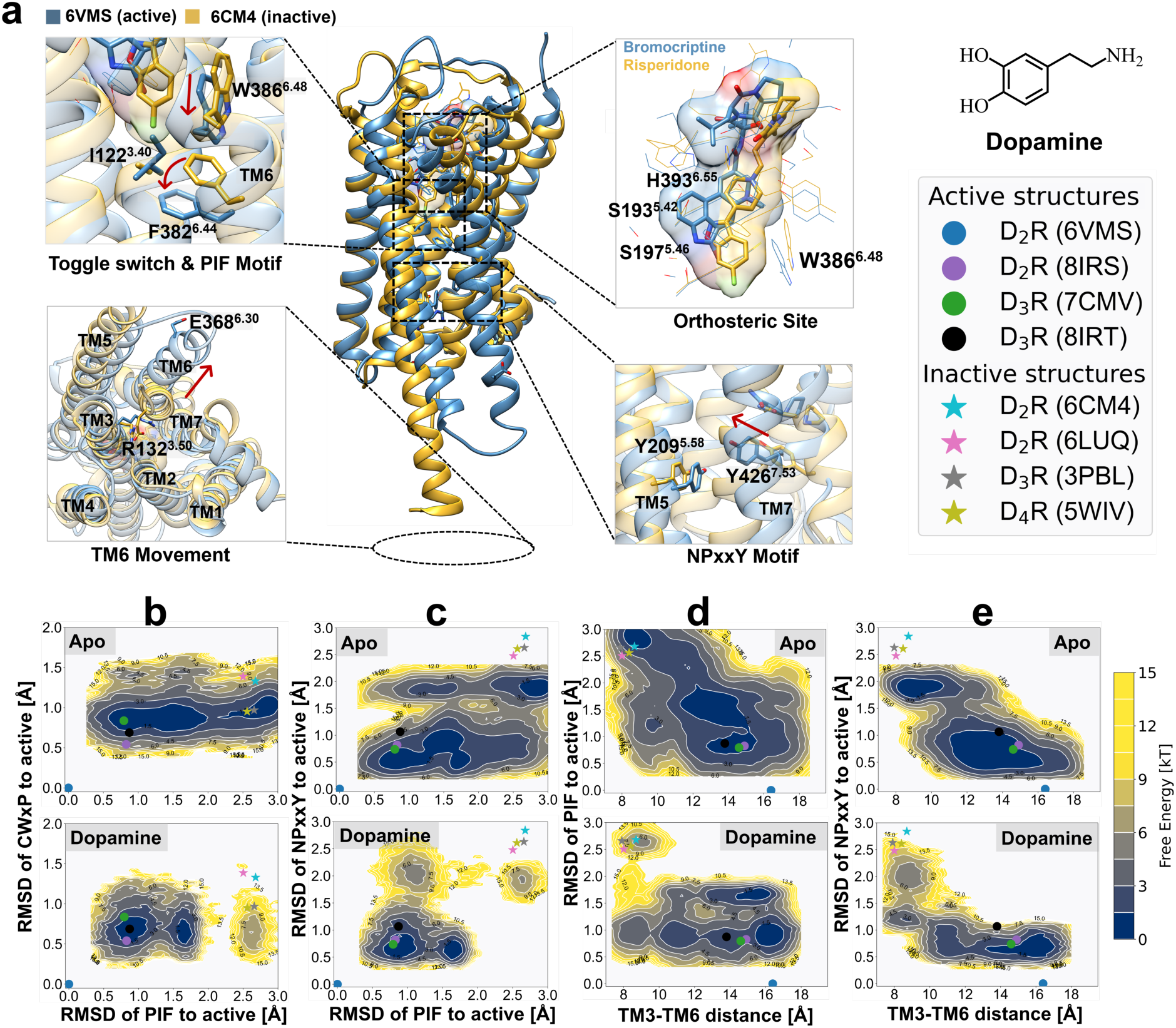
Molecular mechanism of D_2_R activation. (a) Microswitches involved in D_2_R activation. Conformational changes were identified in the toggle switch (W386^6.48^, CWxP motif), connector region (P201^5.50^-I122^3.40^-F382^6.44^), NPxxY motif (N422^7.49^-Y426^7.53^), and intracellular part of TM6. (b-e) Free energy landscapes of apo and dopamine-bound D_2_R projected along (b) the CWxP motif (heavy atom RMSD of C385^6.47^, W386^6.48^, and P388^6.50^), (c) the PIF motif (connector region: heavy atom RMSD of I122^3.40^ and F382^6.44^), (d) the RMSD of the NPxxY motif (heavy atom RMSD of N422^7.49^, P423^7.50^ and Y426^7.53^), and (e) TM6 outward movement (Cα distance between R132^3.50^ and E368^6.30^). In (b-e), the position of representative active and inactive structures of adrenergic and D_2_-like dopamine receptors are depicted by dots and stars, respectively, including active D_2_R (PDB codes: 6VMS and 8IRS), active D_3_R (PDB code: 7CMV and 8IRT), inactive D_2_R (PDB code: 6CM4 and 6LUQ), inactive D_3_R (PDB code: 3PBL), and inactive D_4_R (PDB code: 5WIV).

To characterize the molecular mechanism of D_2_R signaling, the free energy landscapes describing receptor activation in the absence and presence of the endogenous agonist dopamine were calculated using enhanced sampling MD simulations. We first optimized the string method with swarms of trajectories and sampled the conformational transition between the active and inactive states of the D_2_R (17, 25). The string optimization was driven by a set of 33 collective variables (CVs) describing activation. The CVs were derived from equilibrated trajectories of the inactive and active D_2_R using demystifying, a machine learning tool to identify important features from simulations (26). More than 400 iterations of optimization, corresponding to an aggregated simulation time exceeding 2 μs, were carried out for each system to progressively reach the most probable transition pathway that connected the active and inactive conformations with 16 metastable intermediates (Table S1 and Figures S1-S2). The conformational ensembles were then projected onto features representing microswitches (*e.g.*, the RMSD to the active conformation or distances between residues, Table S3), resulting in one- or two-dimensional energy landscapes describing the roles of different motifs in activation (Figure 1 and Figure S16).

Comparisons of the free energy landscapes obtained from the MD simulations revealed the allosteric communication between the orthosteric site and the core of the TM region. Dopamine binding caused a shift in the conformational equilibrium for several of the microswitches. If the free energy landscape was projected onto the RMSD of the CWxP and PIF motifs (Figure 1b), the apo receptor sampled two energy basins representing an inactive state and an intermediate state between active and inactive conformations. Dopamine binding stabilized a more active-like CWxP motif and shifted the equilibrium toward the active PIF conformation. Analysis of MD snapshots showed that dopamine binding pushed W386^6.48^ in the CWxP motif downward and thereby favored an active-like PIF motif.

The energetic coupling between the RMSD of the PIF motif, the RMSD of the NPxxY motif, and the outward movement of TM6 (Cα distance between R132^3.50^ and E368^6.30^) provided insights into the communication between the connector region and G protein binding site (Figure 1c-e). One notable difference observed between the free energy surfaces of the apo and dopamine-bound receptor was that the apo receptor sampled a wider conformational space for all three combinations of microswitches. The landscapes showed that there was dynamic equilibrium between multiple low-energy conformations in the inactive, intermediate, and active-like states. These minima were separated by low energy barriers, suggesting that the apo receptor is intrinsically flexible. Dopamine binding altered the overall energy landscape and typically sampled a subset of the space covered by the apo receptor, favoring active-like states and increasing the barrier to reach the inactive state. The free energy surfaces projected onto the RMSDs of the PIF and the NPxxY motifs showed that dopamine binding preserved the same four energy minima as the apo receptor but shifted the conformational ensemble towards more active-like structures (Figure 1c). The landscapes representing the outward movement of TM6 and RMSD of the PIF motif revealed a loose coupling between the connector region and G protein binding site (Figure 1d). Dopamine binding increased the number of energy basins from three to five, involving three different PIF conformations and TM6 rearrangements in the inactive, intermediate, and active states, respectively. Notably, if the PIF motif adopted an inactive conformation, only the inactive TM6 conformation with the formed ionic lock was observed. However, once the PIF motif was trapped in the active or intermediate conformations, the TM6 transitioned between the inactive, intermediate, and active conformations. These results indicated that the connector region has a critical role in propagating signals to the G protein binding site. We also noticed a dopamine-induced allosteric modulation of the G protein binding site based on the energy landscape representing changes in TM6 and the NPxxY motif (Figure 1e). The dopamine-bound receptor mostly populated fully activate conformations and the inactive state became less favorable compared to the apo receptor. Interestingly, when NPxxY switched between inactive and intermediate conformations, TM6 remained in the inactive state with a formed ionic lock. Only after NPxxY adopted the fully active state, the ionic lock was broken, bringing the receptor to active-like conformations. These findings highlighted the importance of the conformational coordination between the intracellular part of TM6 and TM7 for activation.

Taken together, our simulations suggest that the apo D_2_R is in an equilibrium between inactive and intermediate conformations. The microswitches can undergo independent conformational changes, but the probability of stabilizing certain states of the conserved microswitches are coupled. Upon agonist binding, the activated state is favored and stabilized through the adoption of more active-like conformations of several conserved microswitches.

### Probing the contributions of dopamine functional groups to ligand efficacy

To further evaluate our computational model of D_2_R activation, we performed simulations and pharmacological experiments for 14 dopamine derivatives and serotonin (Figure 2d, Figures S14-15 and Table S4). The potencies (EC_50_) and efficacies (E_max_, relative to the maximal effect of dopamine) were determined by measuring recruitment of mini-Gα_i_ to the human D_2_R using a heterologous cell expression system employing luminescence complementation technology (27, 28). The affinity of each compound for the D_2_R was also determined using radioligand binding assays. These experiments revealed the influence of functional groups on ligand binding and receptor activation (*e.g.* the hydroxyls of the catechol group). Despite their closely related chemical structures, the compounds activated the D_2_R to a different extent, providing a benchmarking set for evaluating the ability of MD simulations to predict ligand activity.

**Figure 2.**
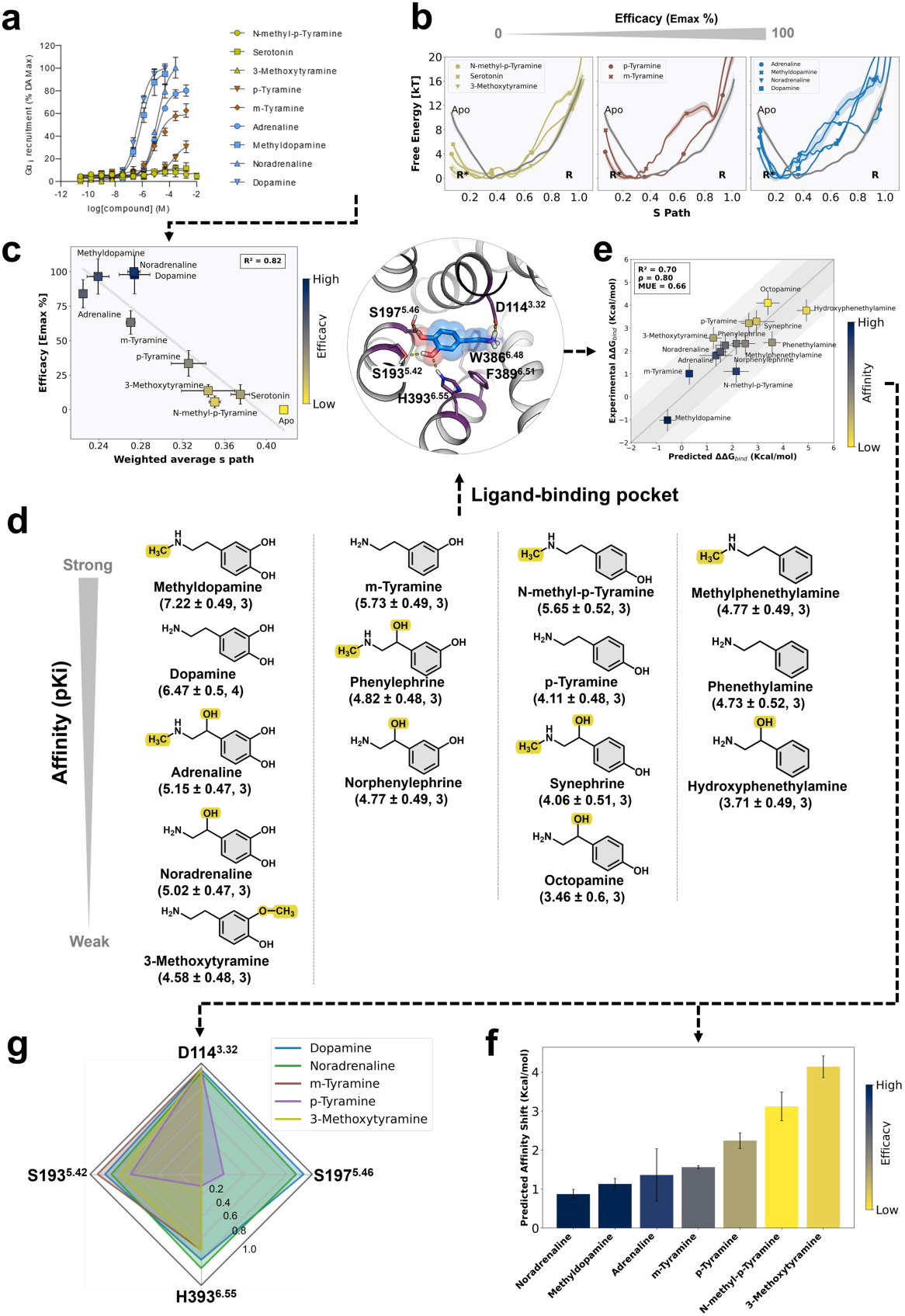
(a) Concentration-response curves for nine compounds tested using a G protein recruitment assay at the human D_2_R. Each point represents the mean ± SEM of three individual experiments consisting of at least duplicates. (b) Free energy landscapes projected along S-path for the different ligands. Weak partial agonists are shown in yellow (N-methyl-p-tyramine, serotonin, 3-methoxytyramine), partial agonists in brown (p- and m-tyramine), and full agonists in blue (adrenaline, methyldopamine, noradrenaline and dopamine). The active and inactive states are marked by R* and R, respectively. (c) Correlation between efficacies of D_2_R ligands and the weighted average S-path values. (d-g) Comparison between calculated and experimental binding free energies of D_2_R ligands. (d) 2D representation of ligands clustered based on chemical structure and sorted by increasing affinity. (e) Correlation between calculated and experimental binding free energies relative to dopamine in the active receptor conformation of the D_2_R. The free energies were calculated from five independent simulations and error bars represent the standard error of the mean (SEM). The coefficient of determination (R^2^), and Spearman’s correlation coefficient (ρ), and the mean unsigned error (MUE, kcal/mol) are shown in the top left corner. (f) Predicted affinity shifts relative to dopamine. The bars are colored based on the efficacy of the ligand, as described in (c). Error bars represent the standard error of the mean (SEM) of five independent FEP calculations. (g) Analysis of receptor-ligand interactions. Radar plots for polar interaction frequency for representative five ligands (dopamine, noradrenaline, m-tyramine, p-tyramine, and 3-methoxytyramine) with four key residues in the binding site (D114^3.32^, S193^5.42^, S197^5.46^ and H393^6.55^).

In order to understand the observed differences in ligand efficacy, enhanced sampling MD simulations of the D_2_R in complex with seven dopamine derivatives and serotonin were performed (Table S1 and Figures S3-10). The dopamine analogs shared a phenethylamine scaffold and were selected to cover the entire efficacy spectrum (Figure 2a). Adrenaline, noradrenaline, and methyldopamine were full agonist whereas modifications involving the hydroxyl groups of the catechol (p- and m-tyramine) resulted in partial agonism. Finally, serotonin and 3-methoxytyramine were ligands of the D_2_R, but did not activate the receptor significantly at the highest tested concentration (Table S4). The other compounds were excluded from these simulations due to low affinity (hydroxyphenethylamine, octopamine, synephrine) or because the E_max_ could not be reliably determined (phenetylamine, norphenylephrine, phenhyleprine, n-methyl-phenetylamine). To characterize the free energy landscape of D_2_R in complex with different ligands, we calculated the S-path, a global feature taking into account overall structural rearrangements that was calculated from all the 33 predefined CVs (Figure 2b) (29). The free energy landscapes were projected along the S-path, which ranges between 0 (fully active conformation) and 1 (inactive conformation). The simulations of ligands with diverse efficacy profiles resulted in differences in the lowest energy basin sampled and energy barriers between states. One common observation across all studied systems was that the receptor tended to sample lowest energy basins with S-path values of less than 0.5. The apo receptor was stabilized in an overall intermediate state with S-path ∼0.4. Weak partial agonists (N-methyl-p-tyramine, 3-Methoxytyramine, and serotonin) led to similar landscapes as the apo receptor but the energy barrier to reach the active state decreased to different extents. Full agonists, such as dopamine and noradrenaline, greatly increased the energy barrier to reach the inactive state and trapped the receptor in a more active-like conformation (S-path = ∼0.2). To quantify the effect of the ligands on the receptor in the simulations, we calculated the weighted average value of S-path for each complex based on the free energy distribution, and compared the results to the experimentally determined efficacies (E_max_ values). Notably, there was a strong linear correlation between weighted average S-path values and ligand efficacies (R^2^ = 0.82, Figure 2c).

To gain additional mechanistic insight, we also explored the relationships between local conformational changes of specific microswitches and the experimentally observed ligand efficacy (Figure 1a). For each ligand, the free energy landscapes were projected along different microswitches, and the correlation between weighted average values and ligand efficacies was evaluated (Figure S16). There was a good correlation between the conformational changes of the CWxP motif and ligand efficacies (R^2^ = 0.71, Figure S16e), which was due to the direct interaction between the ligands and W386^6.48^. The outward movement of TM6 was also correlated with ligand efficacy (R^2^ = 0.81, Figure S16h), reflecting the ligand-induced allosteric modulation of the G protein binding site. However, there was no correlation between the other microswitches that connect the ligand and G protein binding sites (PIF and NPxxY motifs) and ligand efficacies (R^2^ = 0.09 and 0.10, respectively, Figure S16f-g). The differences in efficacy could hence be predicted both by the global structural features captured by S-path and conformational changes of two specific microswitches (CWxP motif and TM6).

The enhanced sampling MD simulations of different compounds were further analyzed to identify conformational states characteristic of full and partial agonism. By comparing basins in the free energy landscape sampled by all studied complexes (Figure 2b and Figure S16), we identified four metastable conformational states along the activation pathway. Four representative snapshots corresponding to these local minima were extracted from the simulation of the dopamine-bound receptor (Figure 1e). The metastable states covered different conformations of the microswitches, including two conformations of the CWxP motif (inactive and active) and three conformations of the PIF motif, NPxxY motif, and TM6 displacement (inactive, intermediate, and active). The snapshot of the inactive receptor conformation, which was similar to the crystal structure of this state, was most populated in the presence of compounds that did not activate the D**_2_**R (N-methyl-p-tyramine, 3-methoxytyramine and serotonin) (Figure 3a). These compounds favored an intermediate inactive-like conformation (Figure 3b), in which the PIF and NPxxY motifs are partially activated, but the ionic lock is formed. Another intermediate conformation (Figure 3c), which was more active-like, was maximally populated by partial agonists such as p-tyramine (Figure S16). In this state, the fully active conformation of the PIF and NPxxY motifs results in the formation of a hydrogen bond between Y209^5.58^ and Y426^7.53^ (YY motif), which facilitates disruption of the ionic lock. This state was also frequently sampled in the simulation with m-tyramine. Compared to p-tyramine, m-tyramine exhibited a higher propensity to populate a fully active state (Figure 3d), in which all studied motifs were active and the receptor had an open G protein binding pocket. This result was consistent with our signaling assays and previously published studies, in which the partial agonist m-tyramine yielded a higher E_max_ value than p-tyramine (30–33). Finally, the active state conformation was the most populated state for the full agonists such as dopamine and methyldopamine (Figure 3d and Figure S16).

**Figure. 3.**
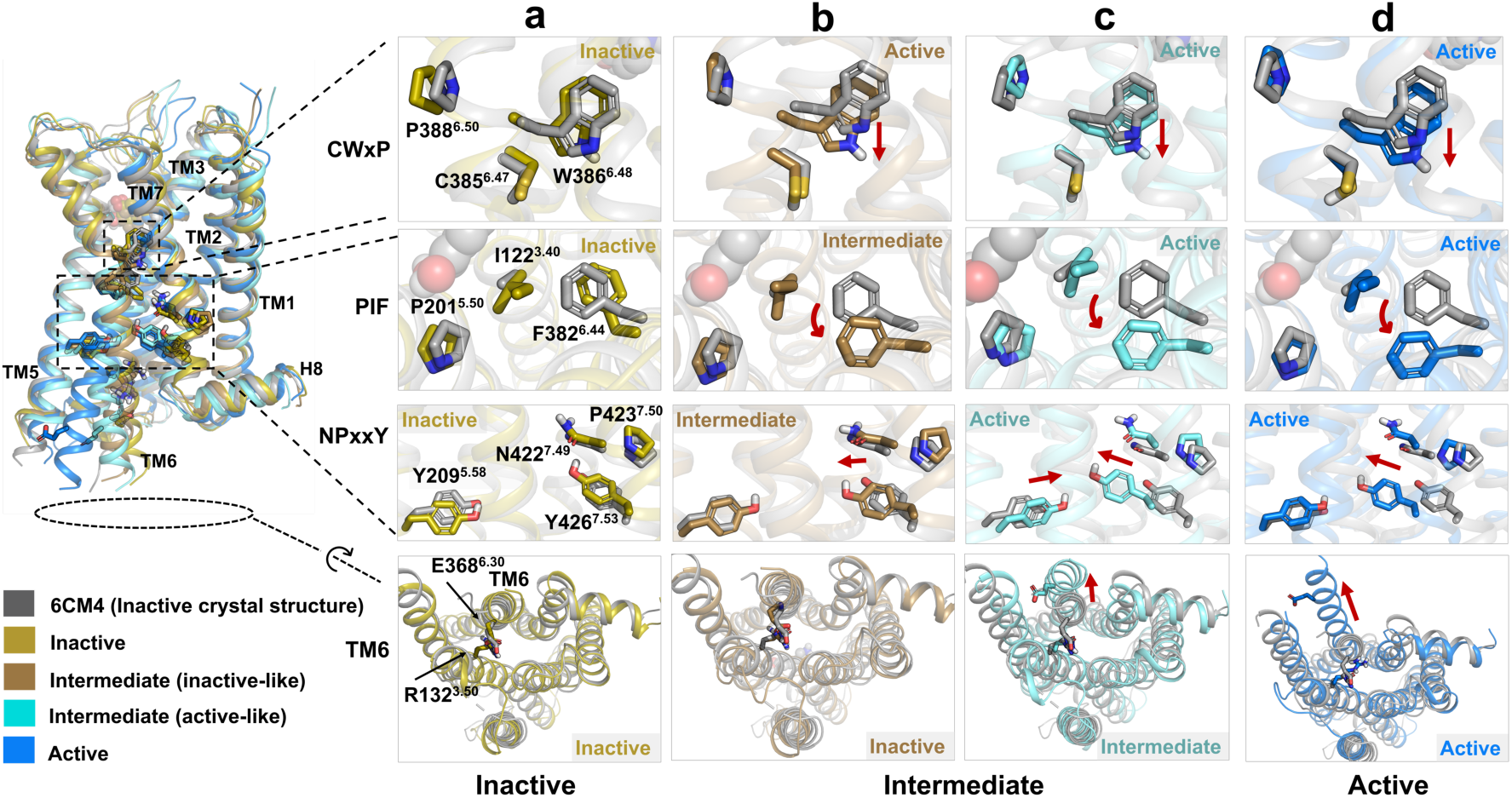
Molecular mechanism of partial and full activation of the D_2_R. Representative simulation snapshots of the dopamine-bound D_2_R in four metastable states describing the conformational transition between the inactive and active conformations: (a) one inactive, (b-c) two intermediate, and (d) one active state. All simulation snapshots were superimposed on the inactive crystal D_2_R structure, which is shown in gray (PDB code: 6CM4).

### Molecular determinants of ligand affinity

To assess if the ligand affinities of the dopamine derivatives could be predicted by MD simulations, we developed protocols for performing free energy perturbation (FEP) calculations. The accuracy of FEP calculations is dependent on access to a high-resolution model of the protein-ligand complex (Figure 2d) (12, 34). Only large ligands (bromocriptine, haloperidol, and risperidone) were bound to the D_2_R in experimentally determined structures and predictions based on these were unlikely to yield good models of the complex with dopamine. We instead generated homology models of the active D_2_R that were primarily based on the cryo-EM structure of the closely related D_3_ subtype in complex with PD-128907 (35), a dopamine derivative. MD simulations of the homology models embedded in a hydrated lipid membrane were performed using spherical boundary conditions. The spherical system was centered on the binding site and was treated as fully flexible, whereas atoms outside a radius of 25 Å were tightly restrained to maintain the receptor in an active conformation. In a first step, FEP calculations of relative binding free energy calculations for dopamine, adrenaline, and noradrenaline were used to identify a model with binding modes that could reproduce the differences in affinity between these ligands. Free energies relative to dopamine were then calculated for all the 14 experimentally evaluated compounds in the series (Figure 2d), which differed in affinity by up to 5800-fold in the radioligand binding experiments. Relative affinities were determined from a perturbation network and the cycle closure error showed that the free energies were well converged (average error = 0.56 kcal/mol, Figure S17). The predicted and experimental relative binding free energies were in good agreement with a mean unsigned error (MUE) of 0.66 kcal/mol, a coefficient of determination (R^2^) of 0.70, and Spearman’s rank correlation coefficient (ρ) of 0.80 (Figure 2e and Table S4). Previous computational studies showed that ligand efficacy can be predicted based on the difference in ligand affinity for the active and inactive states in GPCRs (16, 36, 37). In the case of the D_2_R, the calculated affinity shift values (Figure 2f) also agreed well with the experimental efficacy profiles for the ligands. The three full agonists (methyldopamine, adrenaline, and noradrenaline) had the lowest affinity shifts (average value of 1.1 kcal/mol), the two partial agonists had intermediate values (average value of 3.5 kcal/mol), and the compound with the weakest efficacy had the highest affinity shift values (average value of 6.5 kcal/mol).

Clustering of the compounds based on chemical structure revealed structure-activity relationships for the recognition of dopamine (Figure 2d). Modifications to the catechol group of dopamine had a major impact on affinity. Removing both the hydroxyl groups from the aromatic ring in phenetylamine led to a 55-fold loss of affinity compared to dopamine (pK_i_ = 6.47). The hydroxyl group in the meta-position (m-tyramine, pK_i_ = 5.73) was more important for maintaining affinity than the hydroxyl group in the para-position (p-tyramine, pK_i_ = 4.11). Replacing the meta-hydroxyl of dopamine with a methoxy group also led to a large reduction of affinity (methoxytyramine, pK_i_ = 4.58). Overall similar observations were made for modifications to the catechol groups of adrenaline and noradrenaline, which also contain an additional β-carbon hydroxyl group (adrenaline and noradrenaline) and methyl substituent on the amine group (adrenaline) compared to dopamine. Introducing the β-carbon hydroxyl group in noradrenaline (pK_i_ = 5.02) led to 28-fold weaker affinity at the D_2_R compared to dopamine. To determine the molecular basis of the differences in affinity and efficacy, we analyzed MD simulations of five representative ligands and calculated the frequency of polar interactions with four key residues (S193^5.42^, S197^5.46^, H393^6.55^, and D114^3.32^) in the binding site (Figure 2g). The high-affinity and full agonist dopamine and noradrenaline had polar interactions with all the four residues. All compounds formed a salt bridge with D114^3.32^, but the interactions with the other three residues varied. The compounds m-tyramine and p-tyramine were partial agonists with 14- and 6607-fold lower potencies than dopamine, respectively. These results also agreed well with the difference in affinity (5- and 229-fold, respectively). The more potent and efficacious agonist m-tyramine interacted with both S193^5.42^ and H393^6.55^ to a similar extent as dopamine and adrenaline, but did not interact with S197^5.46^. The weaker agonist p-tyramine also formed a hydrogen bond with S193^5.42^ (60%), but interactions with S197^5.46^ and H393^6.55^ were only observed in 20% and 10% of the simulation snapshots. Finally, the weakest ligand 3-methoxytyramine, which did not activate the D_2_R, interacted with S193^5.42^ and H393^6.55^. However, interactions with H393^6.55^ were formed by the methoxy group, which is a weak hydrogen acceptor compared to the hydroxyl group in dopamine. The most efficacious agonists dopamine, noradrenaline, and m-tyramine were hence able to stabilize the hydrogen bonding network with S193^5.42^ and H393^6.55^ in the active receptor conformation. In the inactive D_2_R receptor conformation, this network cannot be formed due to a different conformation of TM6 that leads to a more open binding site.

### Comparison of dopamine and adrenergic receptor activation mechanisms

Both the D_2_R and β_2_AR belong to the group catecholamine-binding GPCRs and have similar binding sites. Several key residues for ligand recognition are conserved in adrenergic and dopamine receptors, in particular the side chains interacting with the positive charge of the agonist (Asp^3.32^) and the hydroxyls of the catechol (Ser^5.43^ and Ser^5.46^) (8). Our simulations of the ligands suggested that there were distinct differences between the binding modes in the two receptors. Whereas the catechol groups formed similar hydrogen bonds to the serine side chains in TM5, the charged amine group of dopamine was more deeply buried in the D_2_R site (Figure 4a). In the case of the β_2_AR, the crystal structure of the complex with adrenaline showed that the amine group is pushed towards the second extracellular loop due to the β-carbon hydroxyl that is coordinated by Asn^7.49^ and the bulkier Val^3.33^. To assess if these differences influence the molecular mechanism of activation, we also performed enhanced sampling simulations of the β_2_AR and compared the results to the D_2_R.

**Figure 4.**
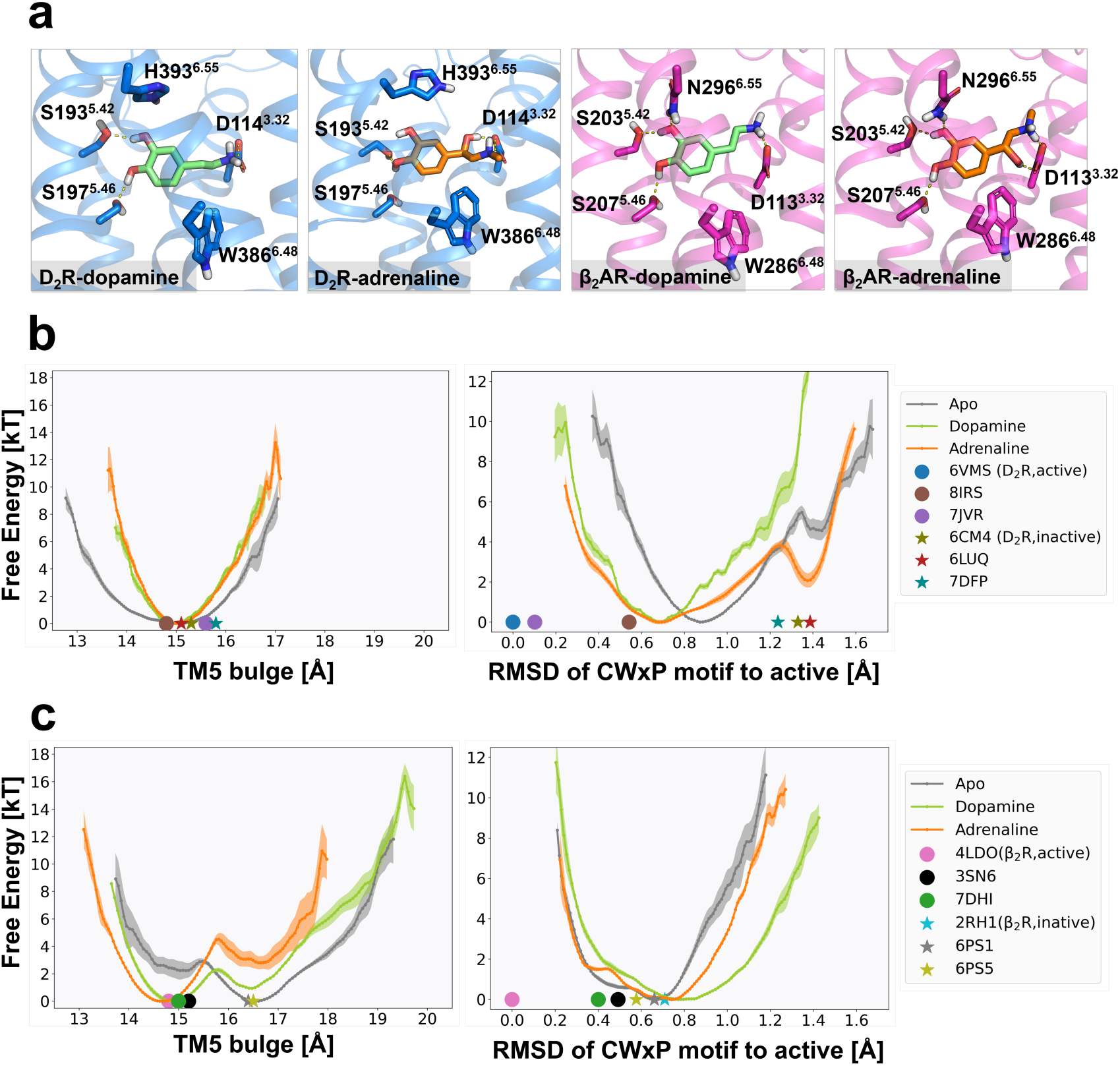
Comparison D_2_R and β_2_AR activation. (a) Binding modes of dopamine and adrenaline in the D_2_R and β_2_AR binding sites from simulations (b-c) Free energy landscapes of D_2_R (b) and β_2_AR (c) with and without agonist bound projected along the TM5 bulge (Cα distance between S^5.46^ and V/M^2.53^) and the CWxP motif relative to the active state (heavy atom RMSD of C^6.47^, W^6.48^ and P^6.50^). Experimental structures of active and inactive states are represented as dots and stars, respectively.

We analyzed four sets of free energy landscapes based on simulations of the D_2_R and β_2_AR in apo forms and bound to their cognate agonist (Figure 4b-c). We focused primarily on the orthosteric site to identify the specific ligand interactions that trigger receptor activation. The free energy landscapes for microswitches in the intracellular region are shown in the Supplementary Information and were overall similar, reflecting that the twist of the NPxxY motif and outward movement of TM6 are conserved features of activation (Figure S18b). Previous studies of adrenergic and dopamine receptors have suggested that conformational changes of Trp^6.48^ in the CWxP motif and interactions with the serine residues in TM5 are involved in the activation mechanism (15, 18, 38–43). We first compared the free energy landscapes obtained for the CWxP motif (Figure 4b-c). In the case of the D_2_R, two minima corresponding to the inactive- and active-like conformations of this motif were identified in the apo simulations. Dopamine binding favored the active state, which primarily reflected a change in the conformation of Trp^6.48^. In contrast, a single free energy basin was stabilized for the apo simulation of the β_2_AR, and adrenaline binding did not influence the Trp^6.48^conformation to the same extent. In TM5, we analyzed the bulge of the helix identified in agonist-bound structures of the β_2_AR, which was described as a consequence of interactions with the two conserved serines (Figure 4c) (44, 45). Analysis of the free energy landscapes of the TM5 bulge identified two minima corresponding to active- and inactive-like states in the simulations of the apo β_2_AR, and adrenaline binding favored the active conformation. However, we observed a single minimum in the free energy landscapes generated from simulations of the D_2_R, and dopamine binding did not change the conformation of the TM5 bulge (Figure 4b). In both the D_2_R and β_2_AR, the observed conformational changes in the CWxP and TM5 bulge, respectively, also favored an active-like PIF motif (Figure 1b and Figure S18a), which connected the orthosteric site to conserved microswitches. These results indicated that the activation mechanisms of the D_2_R and β_2_AR by their cognate agonists were due to different interactions. As dopamine and adrenaline activate both receptors, we performed additional sets of control simulations to assess if the activation mechanism was ligand- or receptor-specific. Simulations of the D_2_R-adrenaline and the β_2_R-dopamine complexes resulted in changes in the free energy landscapes similar to those obtained with the cognate agonist, further supporting a receptor-specific activation mechanism.

## Discussion

The aims of this study were to characterize the activation mechanism of the D_2_R and develop computational approaches to predict ligand affinity and efficacy. MD simulations in combination with free energy calculations and pharmacological assays led to three main results. First, enhanced sampling simulations showed how dopamine shifts the receptor towards active-like conformations by influencing conserved microswitches such as Trp^6.48^ in the binding site. In a second step, we evaluated strategies to profile compounds based on different types of MD simulation techniques, which were able to predict the experimentally determined affinities and efficacies of dopamine derivatives. Finally, comparisons of the D_2_R and β_2_AR showed that these GPCRs are activated by distinct ligand-induced mechanisms despite recognizing closely related compounds. These insights into the molecular basis of activation could accelerate the discovery of novel drugs targeting the dopamine receptors.

A central question for understanding the activation mechanism of GPCRs is how specific interactions in the orthosteric site trigger conformational changes. Previous experimental and modeling studies of the dopamine receptors have suggested that hydrogen bonding with TM5 leads to receptor activation (15, 18, 41, 43). On one hand, this hypothesis is supported by structure-activity relationships of the catechol moiety, which show that the hydroxyl groups were important for achieving full agonism (15, 43). On the other, there are several agonists based on the phenethylamine scaffold (e.g., 2-N,N-dipropylaminotetralin) that do not possess the hydrogen bonds needed to interact with the serine residues in TM5 (32). Our MD simulations supported that D_2_R activation by dopamine is primarily driven by modulation of Trp^6.48^, which is consistent with previous studies demonstrating that this microswitch plays a key role in the activation of several GPCRs (18, 20, 46, 47). In our simulations of the apo state, the D_2_R exists in an equilibrium between inactive, intermediate, and fully active states. Dopamine binding induces a downward shift of Trp^6.48^, which stabilizes more active-like conformations of conserved microswitches such as the PIF and NPxxY motifs. Other binding site interactions may indirectly affect ligand interactions with Trp^6.48^, be important for high affinity binding, or differentially influence activation of different effectors such as G proteins and β arrestins (15). We anticipate that these results can guide discovery of novel dopamine receptor agonist scaffolds.

One of the challenges in GPCR drug discovery is to optimize the efficacy of a ligand for therapeutic effect. Our results show that MD simulations hold promise as a profiling tool to probe how chemical modifications will affect the activity of a compound. As a proof-of-principle, we studied derivatives of the endogenous ligand, a strategy that has been very successful in GPCR drug discovery, and two complementary simulation techniques were explored. We first performed enhanced sampling simulations based on the string of swarms method, which were able to predict the efficacy of a compound from free energy landscapes describing the conformation of the receptor. This technique provides insights into the unique ensemble of receptor conformations stabilized by a ligand, which could potentially enable design of functionally selective agonists (48). However, enhanced sampling simulations require receptor-specific selection of collective variables and are computationally demanding. Consequently, such calculations can only be performed for small sets of compounds. It is also worth noting that the string method simulation is dependent on the predefined initial pathway, and thus it is possible that other transition pathways or activation mechanisms co-exist. The second technique involves calculations of binding free energies in the active and inactive conformations of the receptor, which was recently shown to be effective for diverse GPCRs (16, 36, 37). Such simulations can profile larger sets of compounds and, in contrast to the enhanced sampling approach, also predict relative ligand affinities. However, high-resolution starting structures of receptor-ligand complexes are required to obtain accurate predictions (34), and our approach neglects that there may be multiple distinct active or inactive receptor conformations. After optimizing the structure of the D_2_R-dopamine complex, binding free energy calculations resulted in good predictions of compound affinities and efficacies. In applications to hit-to-lead optimization, we envision a multi-stage approach utilizing the strengths of each technique to prioritize candidates for experimental evaluation.

The major conformational changes upon GPCR activation are conserved in the class A family, which is remarkable considering the vastly different endogenous ligands recognized by the orthosteric sites (4). The dopamine and adrenergic receptors both bind similar catechol agonists, but have evolved to selectively recognize their cognate ligand among a large number of other biogenic amines. For example, adrenaline is a >1000-fold more potent agonist of the β_2_AR than dopamine (49), which is intriguing considering that the binding sites of the adrenergic and dopamine receptors are highly similar. Our simulations indicated that ligand-induced activation of the D_2_R and β_2_AR is not primarily driven by the same interactions, which provides novel insights into the mechanisms of receptor selectivity. Conserved microswitches in the intracellular part of the TM region undergo similar conformational changes (4, 20, 44), but the small side chain re-arrangements in the binding site are different in the D_2_R and β_2_AR. Whereas interactions with Trp^6.48^ are critical for activation of the D_2_R, hydrogen bonding with serine side chains in TM5 appears to be the key interaction in the β_2_AR. These differences are partly due to a difference in the ligand binding mode, which explains structure activity relationships of D_2_R and β_2_AR agonists. Recently determined cryo-EM structures of active D_2_-like dopamine receptors show that the positively charged groups of synthetic agonists (e.g. Rotigotine and PD128907) both form a salt bridge with Asp^3.32^ and extensive van der Waals interactions with Trp^6.48^. The corresponding charged groups of β_2_AR agonists also interact with Asp^3.32^, but are not as deeply buried in the binding site of the receptor and form π-cation interactions with Phe193^EL2^ in the second extracellular loop. In agreement with our results, the characteristic bulge of TM5 in experimental structures of the active β_2_AR has not been observed in the D_2_R structures. Our results illustrate that activation mechanisms identified in one GPCR may not even be transferrable to other closely related receptors. MD simulations can serve as a tool to identify the key ligand interactions in the receptor of interest and guide agonist design.

Access to atomic resolution structures of GPCRs has the potential to accelerate drug design, but the static view of activation provided by experimental methods generally cannot explain how chemical modifications to agonists alter affinity and efficacy. The MD simulation approaches developed in this work can provide unique mechanistic insights that improve understanding of the complex process of receptor activation. In drug discovery applications, the compound profiling tools that can be applied to a large number of targets in the GPCR family.

## Materials and Methods

### String-of-swarms molecular dynamics simulations

#### Simulation system setup

All dopamine receptor systems were prepared with the initial coordinates taken from the active (PDB ID 6VMS (20)) and inactive (PDB ID: 6CM4 (23)) D_2_R crystal structures. Five mutations T205^5.54^I, M374^6.36^L, V381^6.40^Y, V384^6.43^L and V421^7.48^I in 6VMS and three mutations I122^3.40^A, L375^6.37^A, and L379^6.41^A in 6CM4 were reverted. Two histidine residues, H106^3.24^ and H393^6.55^, were protonated at their epsilon position. All adrenergic receptor systems were prepared with initial coordinates from the active (PDB ID 4LDO (44)) and inactive (PDB ID 2RH1 (50)) β_2_ΑR crystal structures. Two histidine residues H93^2.64^ and H296^6.58^ were protonated at their delta and epsilon position, respectively. Each system was prepared using CHARMM-GUI (51) with the CHARMM36m force field (52). The protein was inserted in a POPC membrane bilayer with 140 lipids and solvated into a TIP3P water box (53) with 0.15M concentration of neutralizing sodium and chloride ions. MD simulations were performed with GROMACS 2020.5 in an NPT ensemble at 300 K. The simulation systems were minimized, thermalized, and equilibrated under gradually decreasing constraints from 2000 to 0 kJ/mol/Å based on the default setting on CHARMM-GUI. The Verlet cut-off scheme was used. All bonds involving hydrogen atoms were constrained with the LINCS algorithm (54). The particle mesh Ewald (PME) method (55) was applied to treat the long-range electrostatic interactions. Velocity rescaling with a stochastic term and Parrinello-Rahman settings were used to deal with temperature and pressure coupling respectively. A time step of 2 fs was used. Nine ligands studied were docked into 6VMS using Glide in Schrödinger Suite 2021 (56, 57). All simulation details involved in this study were summarized in Table S1.

#### Collective variable selection

Collective variables (CVs) for enhanced sampling should capture slow degrees of freedom of the system, separate different functional conformations and characterize the overall process of activation. In this study, we used the Demystifying toolbox (26) to derive CVs for enhanced sampling MD simulations. Demystifying applies machine learning and dimensionality reduction methods to identify significant features from simulation trajectories and scales the corresponding importance profiles between 0 and 1. We applied this tool here to classify snapshots extracted from ∼500 ns equilibrated active and inactive D_2_R simulation trajectories, respectively. As features to train a Random Forest model (RF), we chose inverse inter-residue Cα distances with 200 estimators. More than 50 Cα distances were filtered with the importance profiles over 0.50, of which 32 Cα distances were finally selected by visual inspection to serve as CVs. We also included the Cα distance between R^3.50^ and E^3.68^ as a CV to capture the conformational changes at the G protein binding site. The disruption of this salt bridge (“ionic lock”) is a hallmark of class A GPCRs activation (4). In total, 33 CVs were used for the following enhanced sampling simulations (Table S2). The CVs used for β_2_AR simulations were described in previous work (17).

#### String method with swarms of trajectories

The string method with swarms of trajectories aims to iteratively refine an initial transition pathway toward the most probable transition pathway connecting two stable states in a high-dimensional space supported by a set of CVs. The optimization procedure relies on the estimation of the average drift of the CVs from swarms of short unbiased trajectories. Here, we used an optimized version of this method, developed in our previous work (25) and successfully applied to diverse macromolecules including GPCRs (17, 25), sugar transporters (58) and ion channels (59), to investigate the conformational transitions of D_2_R in the presence of various ligands along different microswitches. For each system, a nonequilibrium steered molecular dynamic (SMD) simulation was performed to obtain an initial transition pathway between the active and inactive receptor states. All the simulations were started from the equilibrated initiated active conformation (D_2_R: 6VMS, β_2_AR: 4LDO) in the absence and presence of different ligands. A restraint constant of 10,000 kJnm^−2^ was applied on all CVs (33 in D_2_R and 41 in β_2_AR) mentioned above simultaneously to pull the receptor from active to inactive states. Except for the initial active conformation, 17 snapshots in each SMD simulation were extracted evenly along the pulling pathway at a 6 ns interval. The last snapshot was visually inspected and compared to the inactive crystal structure (D_2_R: 6CM4, β_2_AR: 2RH1) to ensure the receptors had reached an inactive conformation. If the inactive state was not reached, a new SMD simulation with an increased external potential was applied to the corresponding system until the last inactive configuration was reached. After that, an initial transition pathway was generated with 18 points (corresponding to snapshots extracted from SMD simulations) distributed between the active and inactive receptor states.

The string method with swarms of trajectories was performed on the initial pathway with fixed start (active) and end (inactive) points. For each iteration, a 30 ps restrained equilibrium with a 10,000 kJ nm^−2^ harmonic force constant was first launched at each point, followed by swarms of 32 independent 10 ps unbiased trajectories to calculate the average drift of the CV distance. Finally, the string was reparametrized based on the drift at each point. Through repeating the process above, the string was updated iteratively and progressively to reach the most probable transition pathway connecting the active and inactive conformations with multiple metastable intermediates. The simulation details for each system are listed in Table S1 and the corresponding convergence estimates are shown in Figures S1-13.

#### Free energy landscape calculations

A set of one- and two-dimensional free-energy landscapes were calculated along several local/global features via Markov State Model (MSM) with the Deeptime python library (60). A maximum likelihood MSM was used to construct the transition matrix from the swarms of trajectories, in which a time-lagged independent component analysis (tICA) was selected for dimensionality reduction, followed by discretization via k-means clustering. For local features, the RMSDs of several conserved microswitches (CWxP, PIF and NPxxY) as well as distances of the TM5 bulge and ionic lock were computed to build MSMs. The detailed definitions of these features are listed in Table S3. In addition, a global CV describing the global progression of the system along the conformational transition (S-path) was calculated according to Branduardi’s definition (29). This one-dimensional path CV was built combining all 33 CVs ranging from 0 to 1, which represented the active and inactive states, respectively. Given the string convergence of each system, swarms of trajectories from the last 100 to 300 iterations were used for feature extraction. To check the MSM quality, we estimated the uncertainties using a bootstrapping procedure (61), where the new MSMs were constructed and validated based on the corresponding new transition probability matrix. In this work, 100 bootstrapping samples were generated for the standard deviation calculations.

#### Correlation of conformational changes with ligand efficacy

For each ligand, one-dimensional free energy landscapes along the conserved functional features were calculated to illustrate the ligand effect on the conformational selection between the active and inactive D_2_R states. To quantify the effects of ligands on different features selected for MSMs construction, we computed the weighted average of each feature via the equation, 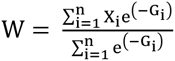, where X_i_ and G_i_ represent the point along the feature and the free energy derived from MSMs, respectively. Based on the weighted average features of ligands and the corresponding E_max_ values (Table S4), the correlation between conformational changes of the receptor and ligand efficacies was computed using linear regression in the scikit-learn package (62). The error for the weighted average features were computed via the equation, 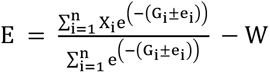, where e_i_ represents the standard deviation from bootstrapping samples.

### Free energy perturbation calculations

#### Generation of starting structures using homology modelling

To generate a model of the active D_2_R-dopamine complex, Modeller v10.2 (63) was thus used to generate 200 receptor structures using a chimera of the active D_3_R structure with PD-128907 (PDB accession code: 7CMV (35)) and β_2_AR active structure with adrenaline bound (PDB accession code: 4LDO (44)). The resulting models were then clustered into 20 representative models using ttClust v4.10 (64) based on the binding site side chains. The resulting 20 models were placed in a 1-palmitoyl-2-oleoyl phosphatidylcholine (POPC) lipid bilayer and equilibrated in GROMACS v2020.4 (65) using the CHARMMGUI protocol (51) with the CHARMM36m (52) force field and TIP3P (53) water model. Protonation states of ionizable residues Asp, Glu, Arg, and Lys in the binding site were set to the most probable in an aqueous solution at pH 7.4, except Asp80^2.50^, which was protonated. Histidine residues in the receptor were protonated at the position δ except for H393^6.55^, which was protonated at the ε position. During equilibration, the backbone of the receptor was held rigid using positional restraints on the heavy atoms whereas the side chains, membrane, and water molecules were allowed to relax. The last snapshot of each of the 20 equilibrated systems was used as starting points for free energy calculations. In order to identify an active model of the D_2_R that could be used in relative binding free energy calculations, we calculated the relative binding free energy of three compounds (dopamine, noradrenaline, and adrenaline) and selected the model with the best prediction of experimental affinities along with low cycle closure error. This receptor model was then used to calculate the relative binding free energies of the remaining ligands. The D_2_R-dopamine in an inactive receptor conformation was generated using Modeller v10.2 based on the inactive D_3_R structure with eticlopride (PDB accession code: 3PBL (66)). A total of 200 models were generated and the top-ranked model was inserted into a POPC membrane. The system was then equilibrated following the same procedures used for the active receptor models. Compounds were either aligned to the dopamine binding mode in the active structure of the D_1_ dopamine receptor (PDB accession code: 7LJD) (67) or to a predicted binding mode of dopamine in the inactive D_2_R generated by docking using Glide (57). Then the Lennard-Jones parameters were generated with OPLS2005 using the ffld_server (68) whereas the partial charges were generated using LigParGen server (69).

#### Free energy perturbation protocol

The FEP simulations were performed with Q (70) using the OPLS all-atom force field version available for this program. The simulations were performed under spherical boundary conditions (SBC) with a sphere radius of 25 Å centered on the ligand. Atoms outside the sphere were excluded from non-bonded interactions. Ionizable residues close to the sphere edge were set to their neutral form and atoms within 5 Å of the sphere edge were restrained to their initial coordinates. Water molecules at the sphere border were subject to radial and polarization restraints according to the surface-constrained all-atom solvent (SCAAS) model (71). Solvent bonds and angles were constrained using the SHAKE algorithm (72). A cutoff of 10 Å was used for non-bonded interactions except for ligand atoms, for which no cutoff was applied. Long-range electrostatic interactions were treated using the local reaction field approximation. In all simulations, a time step of 1 fs was used and non-bonded pair lists were updated every 25 steps. Binding free energies were calculated by transforming one ligand into another in the receptor and in aqueous solution using a single topology protocol that gradually transforms one ligand into another in a series of 93 intermediate states with five replicas per window. The transformations were divided into three major steps: (A) Transformation of partial charges, (B) introduction of soft-core potentials on atoms to be annihilated, (C) transformation of Lennard-Jones potentials and bonded terms. The three steps were typically performed with 31, 11, 51 intermediate states, respectively. At each intermediate state, the receptor-ligand complexes were first equilibrated for 250 ps. During the equilibration, harmonic positional restraints on solute-heavy atoms were gradually released and the system was heated to the target temperature (298 K). This was followed by a 250 ps production phase and energy was collected every 25 fs. The Qfep software (70) was used to calculate the free energy change using Bennet’s acceptance ratio (BAR) (73). Then calculated relative binding free energies were then collected and analyzed using in-house scripts to obtain the mean, the standard error of the mean (SEM), mean unsigned error (MUE), the correlation coefficient (R^2^), and the cycle closure error. To obtain the affinity shift values, alchemical transformations of ligands were performed in the active and inactive states of the receptor relative to dopamine then the free energy in the active state was subtracted from free energy in the inactive state, as described previously (16). To characterize hydrogen bond interactions for the ligands with four key residues in the binding site S193^5.42^, S197^5.46^, H393^6.55^, and D114^3.32^, we used the MDAnalysis package (version 1.1.1) (74) to analyze the five 20 ns trajectories.

### Experimental methods

#### Cell culture and transfections

Expi293F cells were a kind gift from Dr. Ilana Kotliar (Rockefeller University). Cells were maintained in 37 °C, 8% CO_2_ and 95% humidity with continuous orbital shaking, and grown in 125mL polycarbonate Erlenmeyer shake flasks (NEST) using Gibco Expi293 Expression Medium.

#### Competitive radioligand binding assay

To generate D_2_R membranes for radioligand binding assays, cells were expanded to 3e6 cells/mL in 400mL of media in a 2 L polycarbonate Erlenmeyer shake flask (Corning). They were then transfected with a pcDNA.31+ plasmid encoding an N-terminally HA tagged human D2R with a C-terminal -SmBiT tag, using a gene dose of 1μg/mL and ExpiFectamine as a transfection reagent. Transfection enhancers were added the next day and cells collected 48 h post transfection. Collected cells were spun down (150 g for 10 min), washed with dPBS and spun down again. The resulting pellet was re-suspended using a hypotonic lysis buffer (50 mM HEPES, 1% BSA, pH 7.4) for 30 min on ice, then spun down at 20.000 g for 20 min at 4°C. After discarding the supernatant the pellets containing the cell membrane were flash frozen and stored at −80. On the day of each binding assay, the pellets were thawed, re-suspended in membrane resuspension buffer (Tris 50 mM, MgCl2 10 mM, EDTA 0.1 mM, pH 7.4) followed by protein concentration determination using a Pierce BCA protein assay (Thermo Fisher Scientific, Waltham, MA, USA). 30 μg of membrane were incubated, in the dark, for 2 h at RT, in a final volume of 200uL containing [3H]-raclopride (1.5 nM final concentration, PerkinElmer Inc. Boston, MA, USA) and cold ligand (varying concentration) in binding buffer (Tris 50 mM, MgCl2 10 mM, EDTA 0.1 mM, 0.1% BSA, ascorbic acid 100 µM, pH 7.4) similarly as described elsewhere (75), per well in a 96-well plate. Plates were then harvested by vacuum filtration onto 96-well GF type B filter mats (PerkinElmer), pre-soaked with 0.3% polyethylenimine (Sigma, St Louis, MO, USA), using a 96-well harvester (Tomtec, Hamden, CT, USA) and washed three times with cold wash buffer (Tris 50 mM, MgCl2 10 mM, pH 7.4). The filter mats were then dried at 40°C and placed in a sealable bag with 6 mL of Betaplate Scint scintillation cocktail (PerkinElmer). Radioactivity was detected using a Wallac 1450 MicroBeta Trilux scintillation counter (PerkinElmer), using a 3 min tritium protocol.

#### G-protein recruitment assay

G-protein recruitment was assayed using NanoBiT technology as described earlier (27, 28). Briefly, 25 mL of cells (3e6/mL) in 125 mL polycarbonate Erlenmeyer shake flasks were transfected with a 10:1 ratio of HA-hDRD2-SmBiT and LgBiT-mini Gαi for 48h with a gene dose of 1μg/mL. Cells were then diluted 1:4 and resuspended in dPBS supplemented with 10μM coelenterazine-h (NanoLight) with 90μL of cell-suspension added per well to a white 96-well plate (Corning). Luminescence counts (100 ms read time, 100 ms integration time) were measured using a plate-reader (SPARK, Tecan) for 5min to establish a steady baseline before addition of ligands. After ligand addition, luminescence counts were measured for 5min and the collected data was used for subsequent analysis.

#### Statistics and analysis

For radioligand binding experiments, counts per minute (CPM) from the scintillation counter were converted to fM [3H]-raclopride/mg protein per analyzed well, and normalized to the lowest concentration of cold-ligand tested (1e-11M) as 100%. Data were analyzed using “One site - Fit Ki” in Prism 9.5.1 (GraphPad Software Inc., San Diego, CA, USA). For the G protein recruitment assays, luminescence counts underwent background reduction and normalization to a concentration of dopamine evoking a maximal response in the assay system. Data were analyzed and plotted using a preferred model from a comparison of fits between three and four-parameter log(agonist) vs. response function in GraphPad Prism 9.5.1. Data are reported as mean ± standard error of the mean (SEM) with the number of biological and technical replicates indicated in the figure legends.

## Acknowledgements

This work was funded by the Knut and Alice Wallenberg foundation (KAW 2019.0130: P.S., L.D., and J.C.), the Swedish Research Council (2021-4186: J.C.), and the Swedish Strategic Research Program eSSENCE (J.C.). The MD simulations were performed using resources provided by the National Academic Infrastructure for Supercomputing in Sweden (NAISS), partially funded by the Swedish Research Council through grant agreement no. 2022-06725. We acknowledge NAISS, Sweden for awarding this project access to the LUMI supercomputer, owned by the EuroHPC Joint Undertaking hosted by CSC (Finland) and the LUMI consortium. We acknowledge PRACE for awarding us access to Piz Daint, at the Swiss National Supercomputing Centre (CSCS), Switzerland.

