## Supporting Information for "Revealing molecular determinants of ligand efficacy and affinity at the D_2_ dopamine receptor through molecular dynamics simulations"

**This PDF file includes:**

Figures S1 to S18

Tables S1 to S4

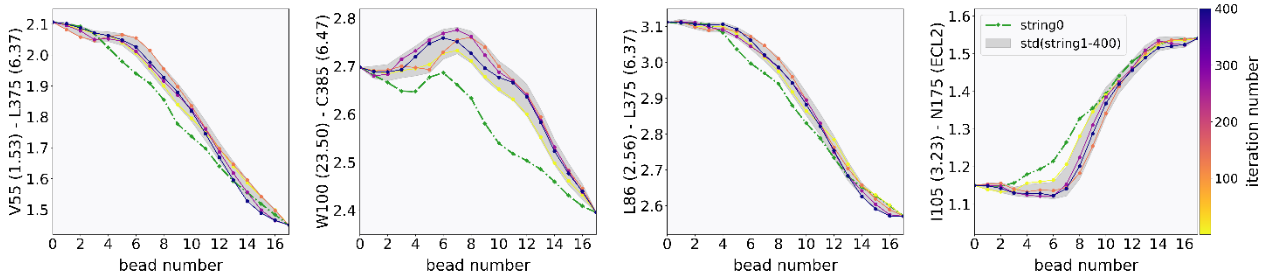


**Fig. S1.** Strings averaged over hundreds of iterations for unliganded D$_2$R initiated from the active structure (PDB accession code: 6VMS). The top four important collective variables (CVs) in Table S1 are shown on the y-axis to evaluate the string convergence. The x-axis shows the evolution of the string points towards the inactive state.

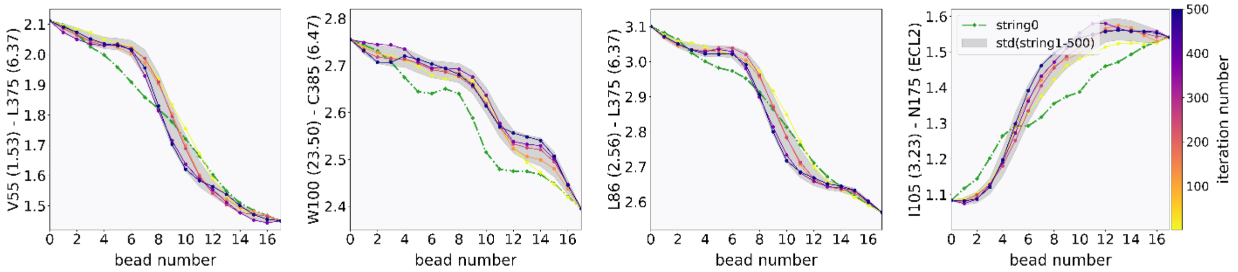


**Fig. S2.** Strings averaged over hundreds of iterations for dopamine-bound D$_2$R initiated from the active structure (PDB accession code: 6VMS). The top four important collective variables (CVs) in Table S1 are shown on the y-axis to evaluate the string convergence. The x-axis shows the evolution of the string points towards the inactive state.

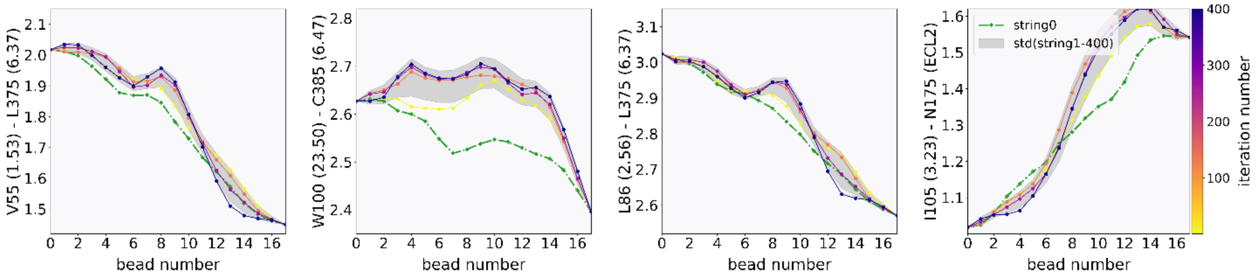


**Fig. S3.** Strings averaged over hundreds of iterations for N-methyltyramine-bound D$_2$R initiated from the active structure (PDB accession code: 6VMS). The top four important collective variables (CVs) in Table S1 are shown on the y-axis to evaluate the string convergence. The x-axis shows the evolution of the string points towards the inactive state.

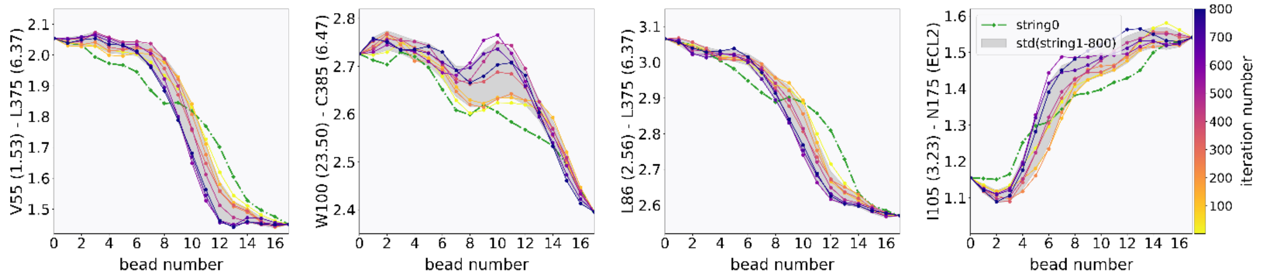


**Fig. S4.** Strings averaged over hundreds of iterations for 3-Methoxytyramine-bound $D_2R$ initiated from the active structure (PDB accession code: 6VMS). The top four important collective variables (CVs) in Table S1 are shown on the y-axis to evaluate the string convergence. The x-axis shows the evolution of the string points towards the inactive state.

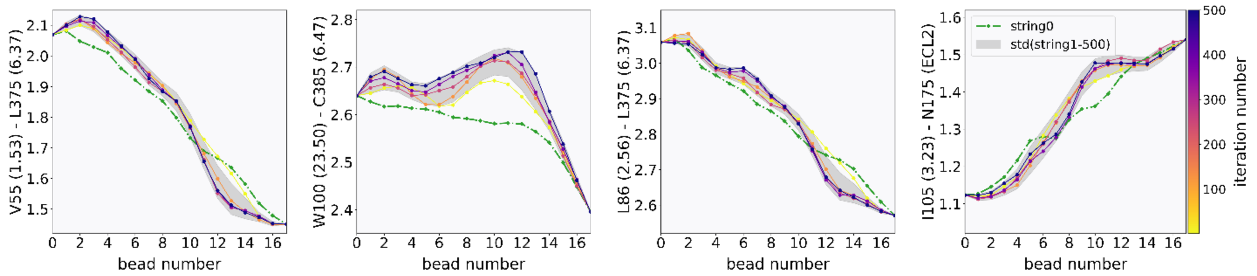


**Fig. S5.** Strings averaged over hundreds of iterations for serotonin-bound $D_2R$ initiated from the active structure (PDB accession code: 6VMS). The top four important collective variables (CVs) in Table S1 are shown on the y-axis to evaluate the string convergence. The x-axis shows the evolution of the string points towards the inactive state.

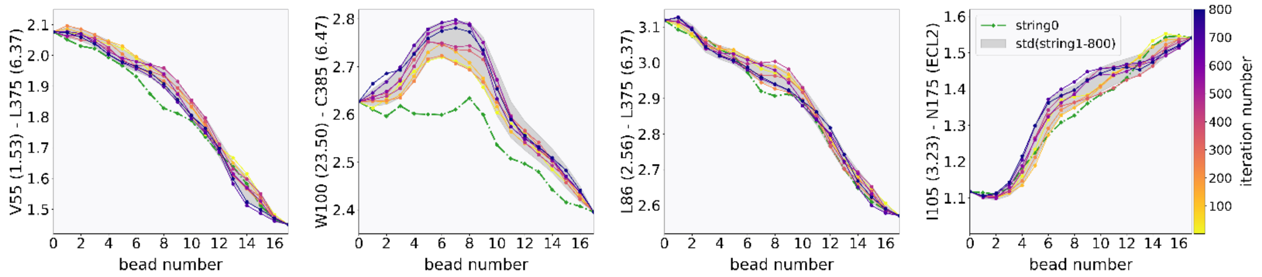


**Fig. S6.** Strings averaged over hundreds of iterations for p-tyramine-bound $D_2R$ initiated from the active structure (PDB accession code: 6VMS). The top four important collective variables (CVs) in Table S1 are shown on the y-axis to evaluate the string convergence. The x-axis shows the evolution of the string points towards the inactive state.

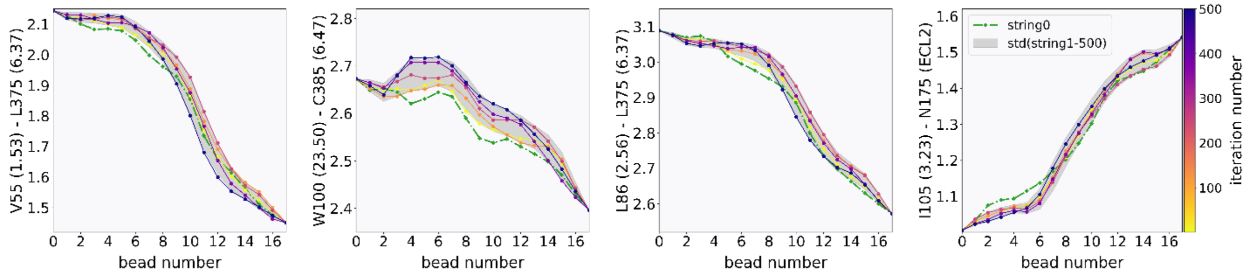


**Fig. S7.** Strings averaged over hundreds of iterations for m-tyramine-bound $D_2R$ initiated from the active structure (PDB accession code: 6VMS). The top four important collective variables (CVs) in Table S1 are shown on the y-axis to evaluate the string convergence. The x-axis shows the evolution of the string points towards the inactive state.

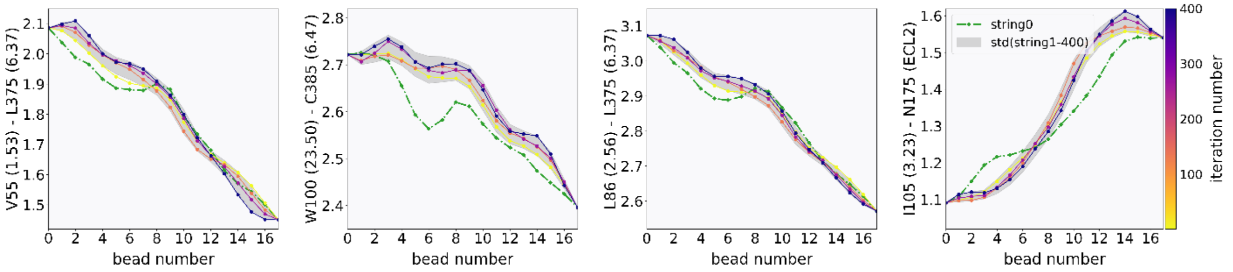


**Fig. S8.** Strings averaged over hundreds of iterations for adrenaline-bound $D_2R$ initiated from the active structure (PDB accession code: 6VMS). The top four important collective variables (CVs) in Table S1 are shown on the y-axis to evaluate the string convergence. The x-axis shows the evolution of the string points towards the inactive state.

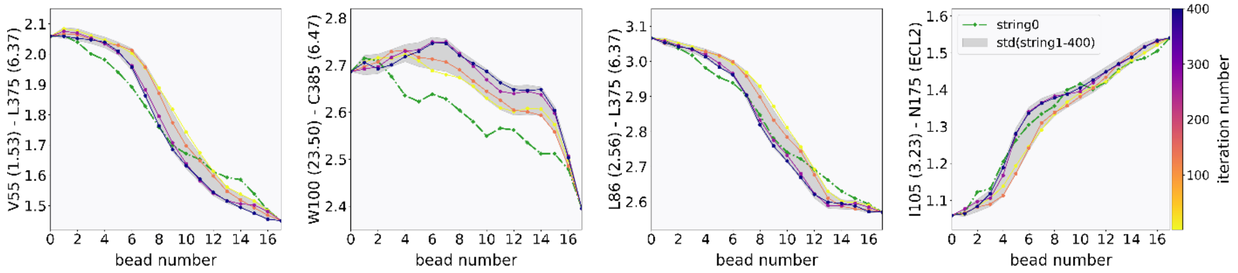


**Fig. S9.** Strings averaged over hundreds of iterations for N-methyldopamine-bound $D_2R$ initiated from the active structure (PDB accession code: 6VMS). The top four important collective variables (CVs) in Table S1 are shown on the y-axis to evaluate the string convergence. The x-axis shows the evolution of the string points towards the inactive state.

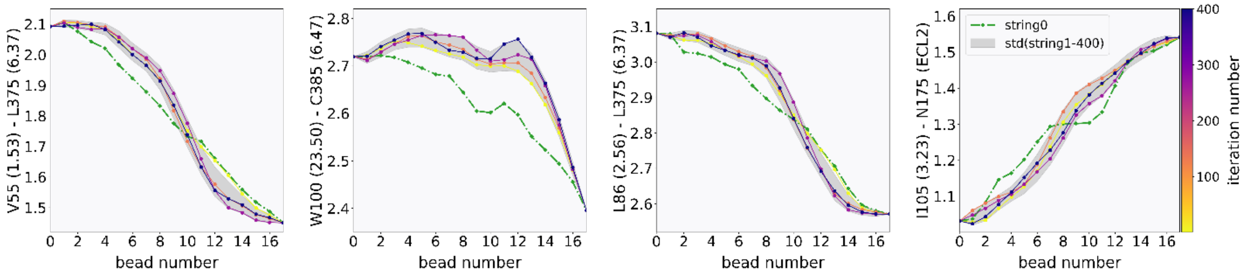


**Fig. S10.** Strings averaged over hundreds of iterations for noradrenaline-bound $D_2R$ initiated from the active structure (PDB accession code: 6VMS). The top four important collective variables (CVs) in Table S1 are shown on the y-axis to evaluate the string convergence. The x-axis shows the evolution of the string points towards the inactive state.

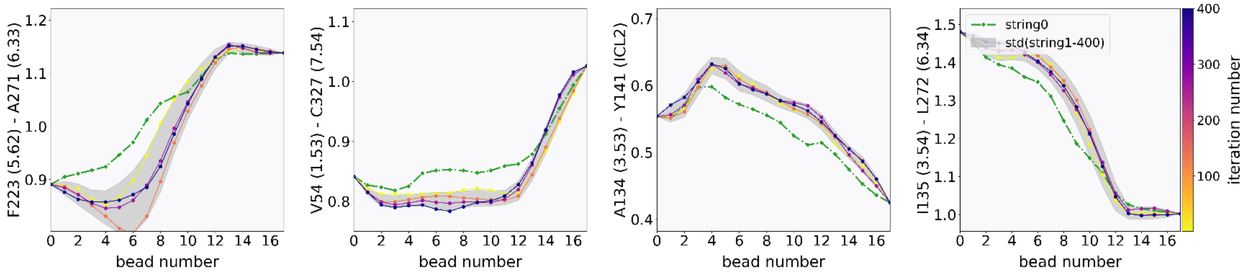


**Fig. S11.** Strings averaged over hundreds of iterations for unliganded $\beta_2$AR initiated from the active structure (PDB accession code: 4LDO). Four important collective variables (CVs) are shown on the y-axis to evaluate the string convergence. The x-axis shows the evolution of the string points towards the inactive state.

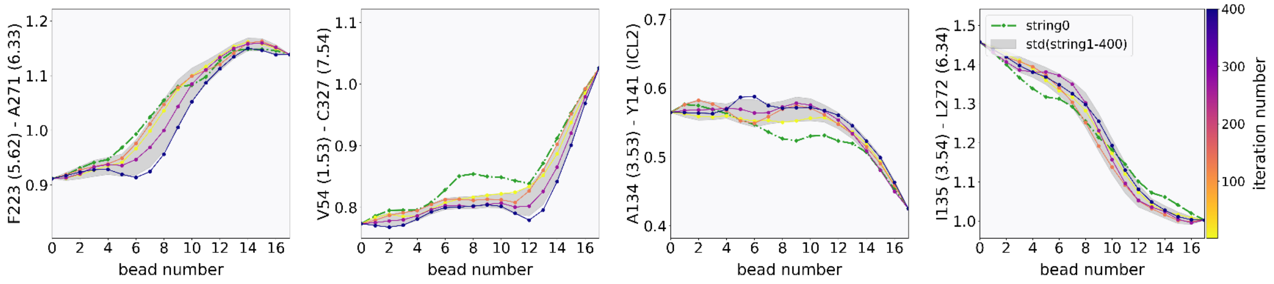


**Fig. S12.** Strings averaged over hundreds of iterations for adrenaline-bound $\beta_2$AR initiated from the active structure (PDB accession code: 4LDO). Four important collective variables (CVs) are shown on the y-axis to evaluate the string convergence. The x-axis shows the evolution of the string points towards the inactive state.

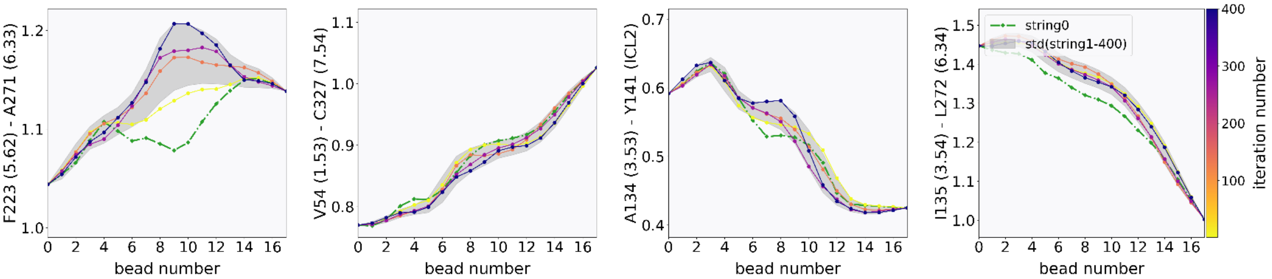


**Fig. S13.** Strings averaged over hundreds of iterations for dopamine-bound β2AR initiated from the active structure (PDB accession code: 4LDO). Four important collective variables (CVs) are shown on the y-axis to evaluate the string convergence. The x-axis shows the evolution of the string points towards the inactive state.

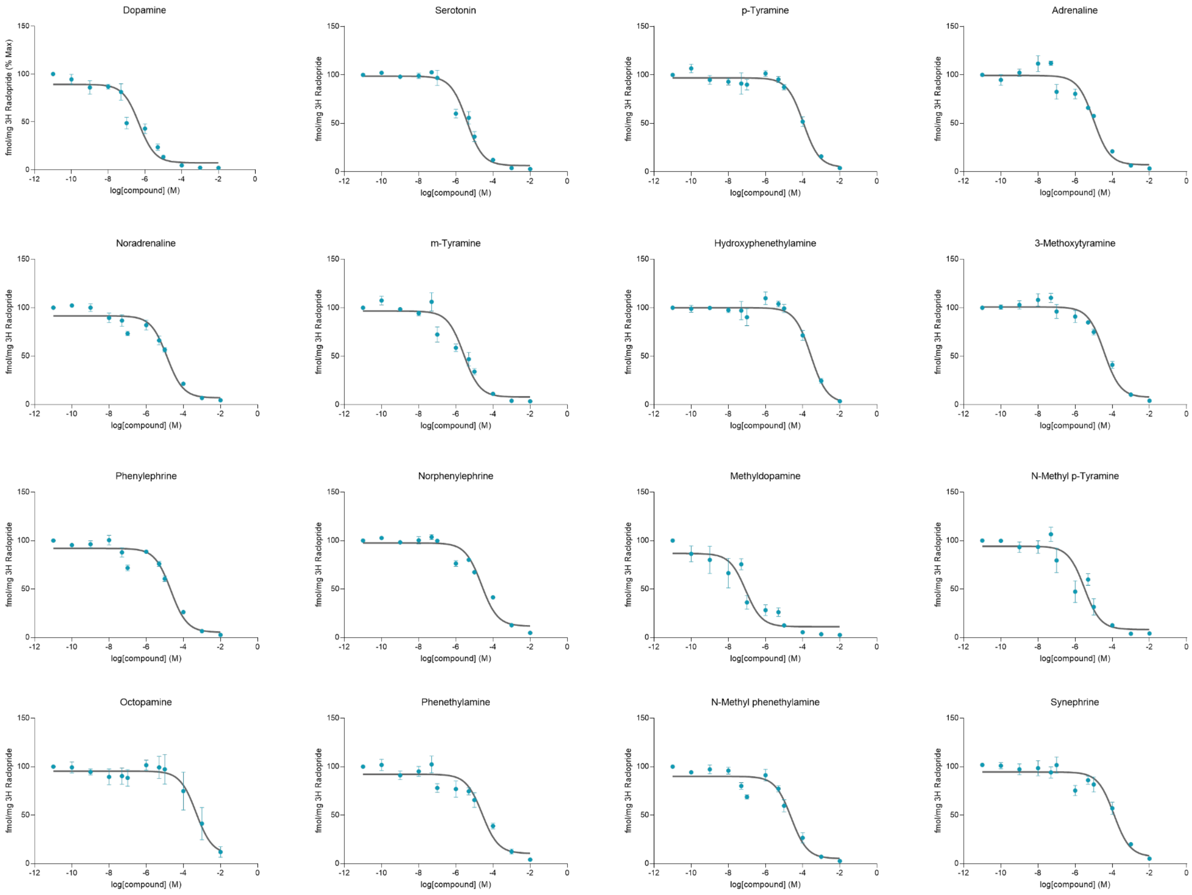


**Fig. S14.** Competitive radioligand binding assay curves for each ligand tested at the human $D_2R$. Each point represents the mean ± SEM of three to four individual experiments consisting of duplicates.

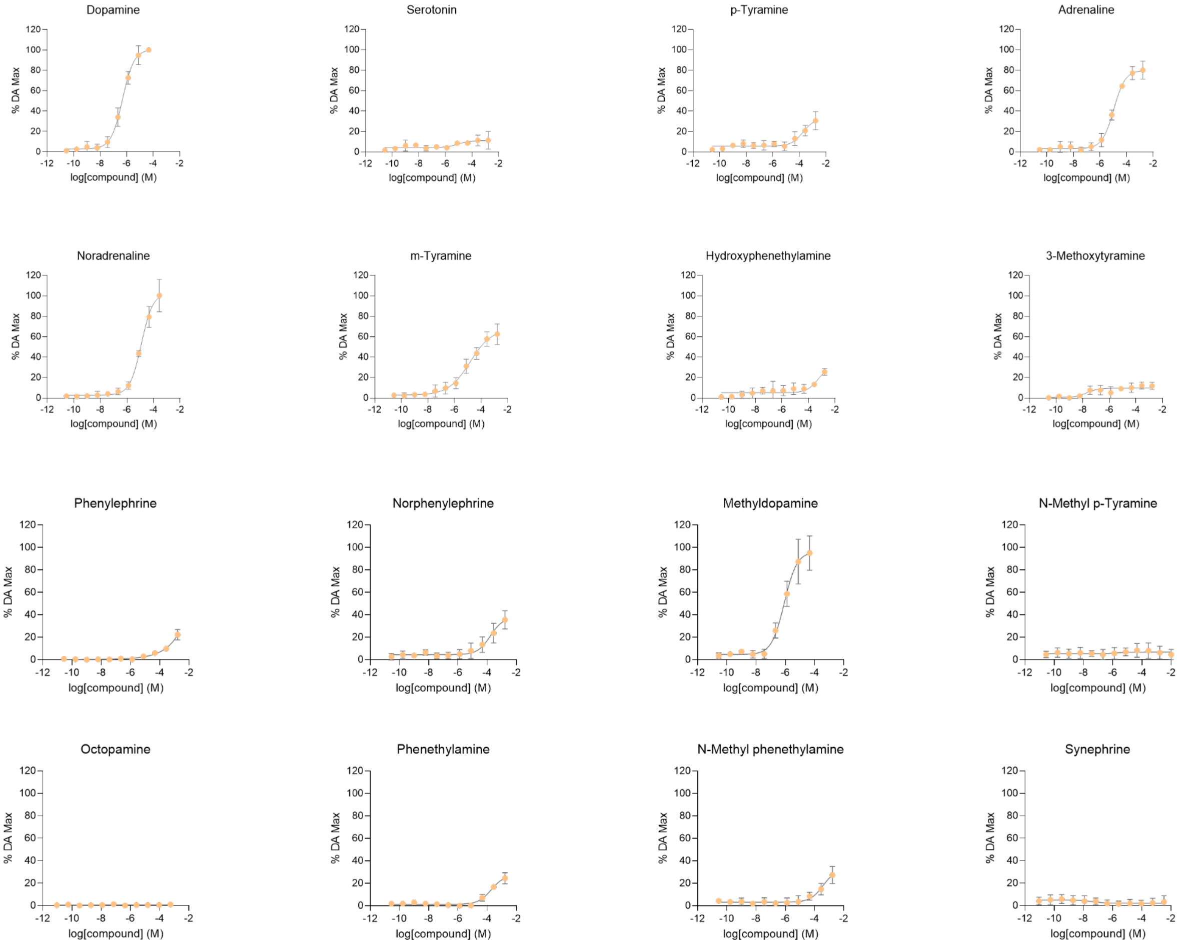


**Fig. S15.** Concentration-response curves for each compound were tested using the G protein recruitment assay at the human $D_2R$. Gαi-recruitment was inferred from luminescence counts resulting from NanoBiT complementation of SmBiT tagged $hD_2R$ and LgBiT tagged mini-Gαi. Each point represents the mean ± SEM of three individual experiments consisting of at least duplicates.

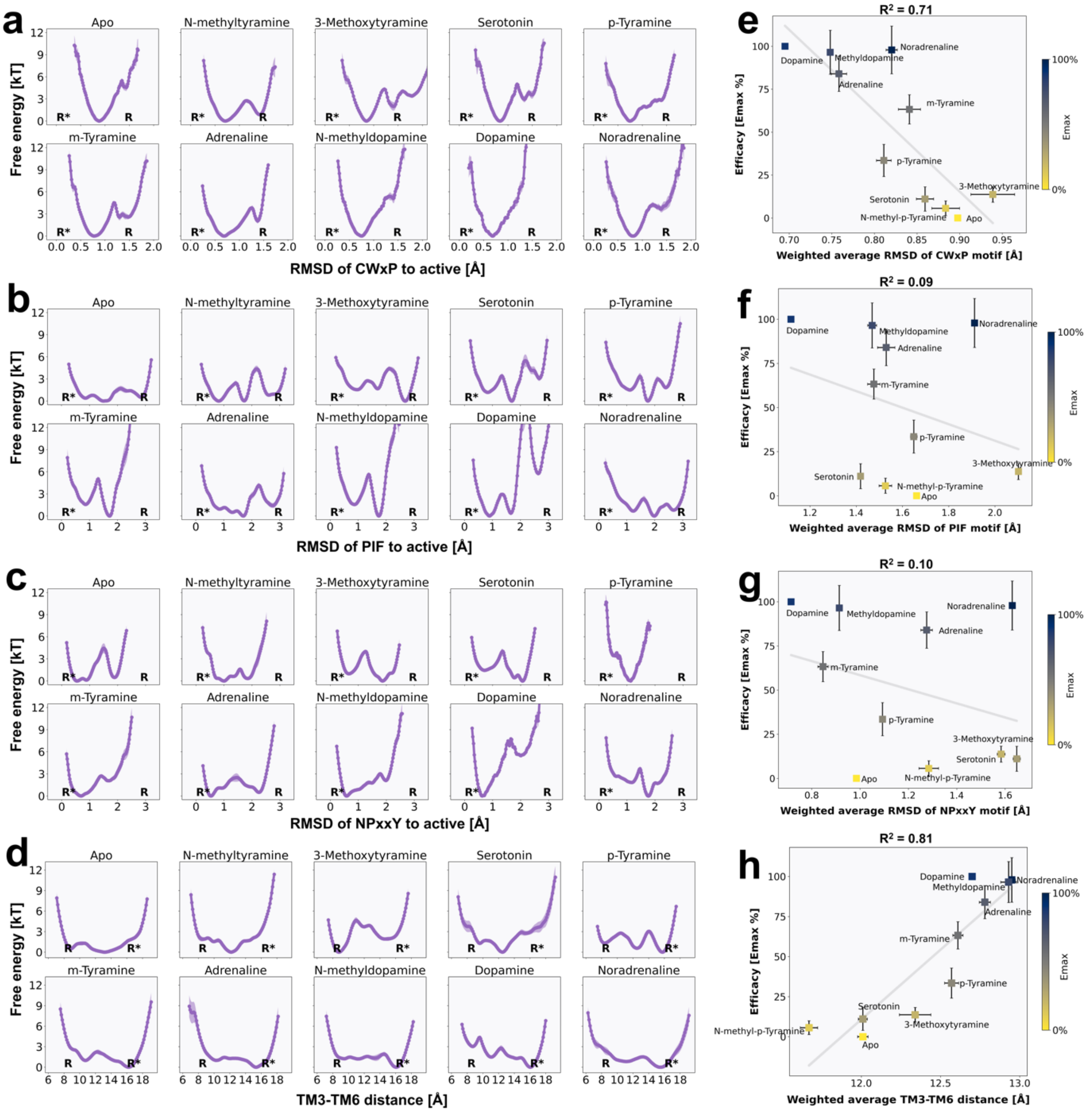


**Fig. S16.** (a-d) Free energy landscapes of $D_2R$ in complex with different ligands projected along a) RMSD of CWxP motif, b) RMSD of PIF motif, c) RMSD of NPxxY motif and d) the outward movement of TM6. The active and inactive states are marked by R* and R respectively. (e-h) Correlation between efficacy and the weighted average of e) RMSD of CWxP motif, f) RMSD of PIF motif, g) RMSD of NPxxY motif, and h) the outward movement of TM6.

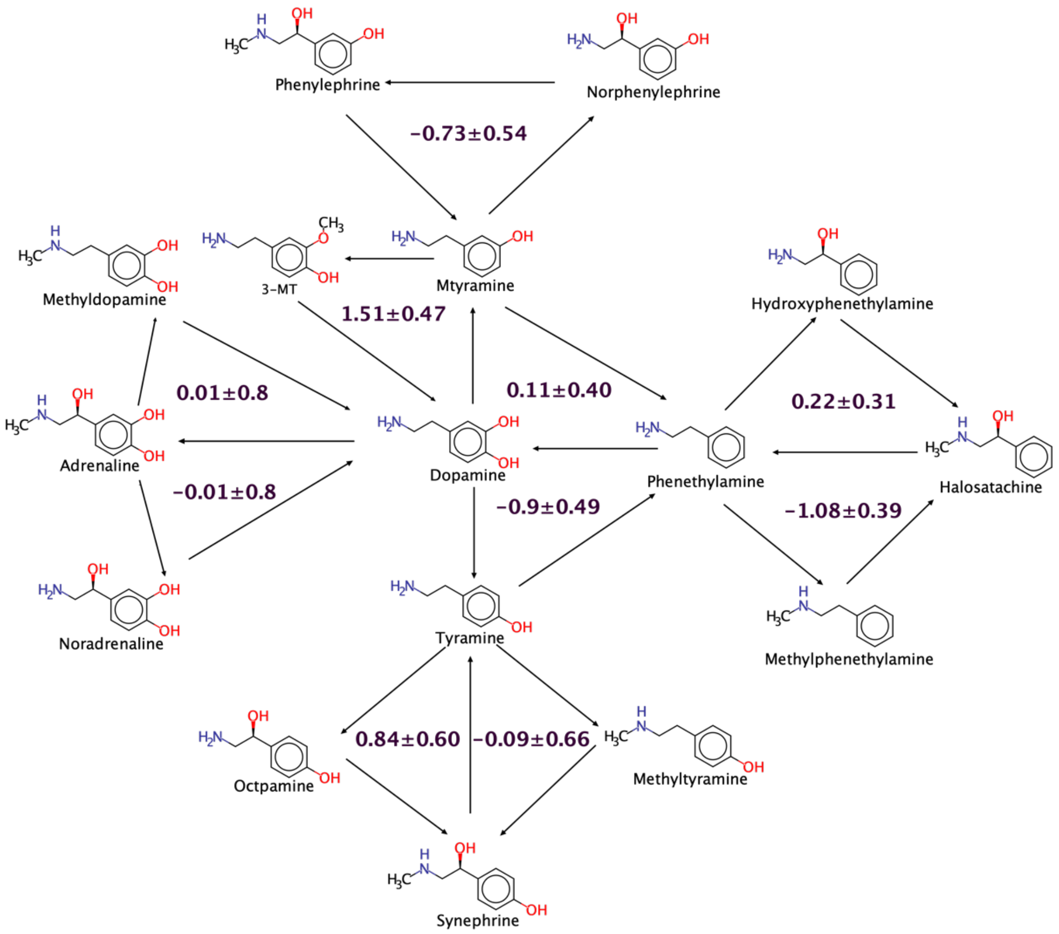


**Fig S17.** Perturbation network used to calculate binding free energies (kcal/mol) relative to dopamine in $D_2R$ active conformation, the cycle closure error is shown in the center of each thermodynamic cycle.

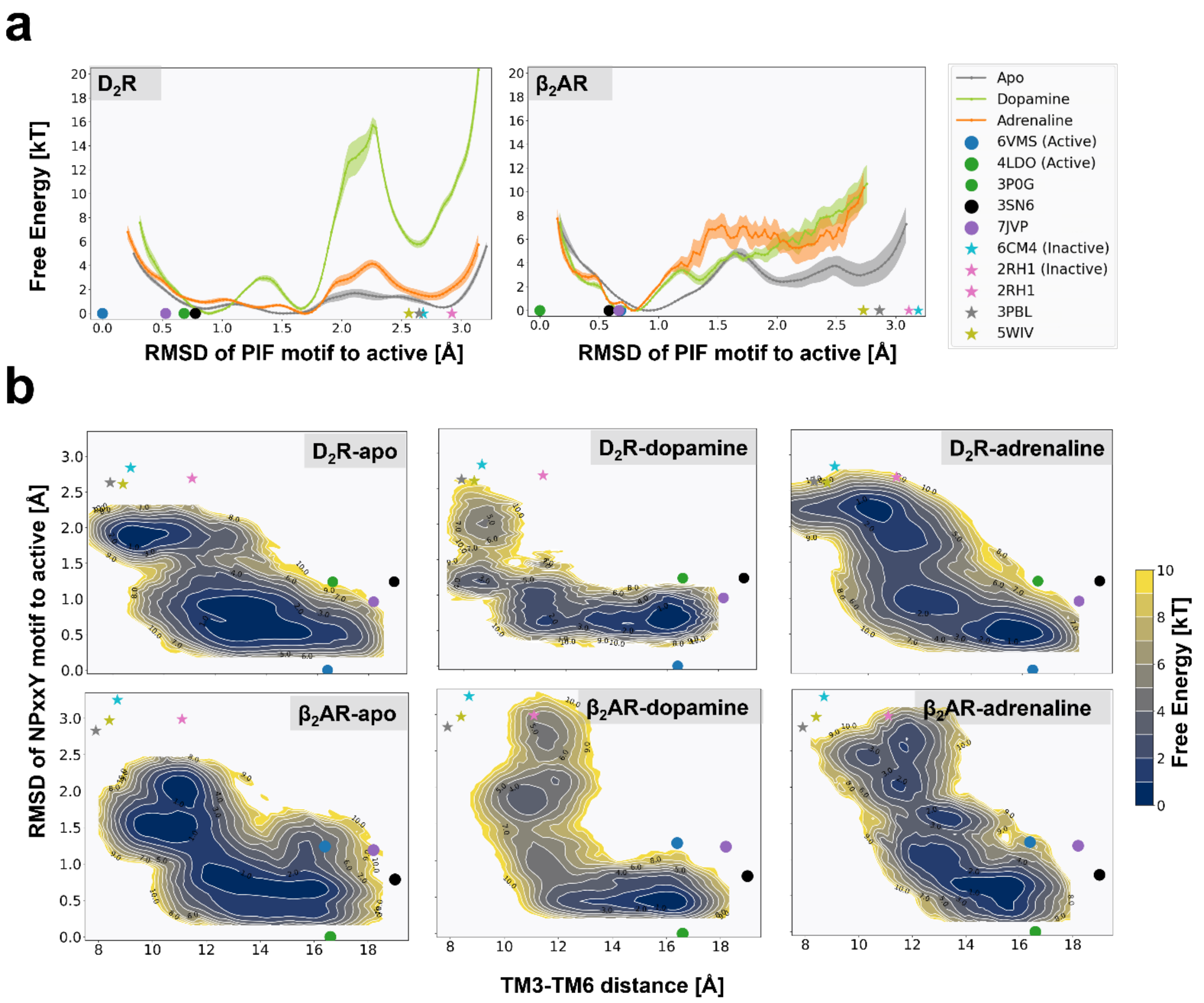


**Fig S18**. Free energy landscapes of $D_2R$/$\beta_2AR$ with and without dopamine/adrenaline bound projected along a) the RMSD of PIF motif relative to the active state (measured on the heavy atoms of $I^{3.40}$ and $F^{6.44}$), and b) the outward movement of TM6 (measured as the distance between Cα atoms of $R^{3.50}$ and $E^{6.30}$) and the RMSD of NPxxY motif relative to the active state (measured on heavy atoms $N^{7.49}$, $P^{7.50}$, and $Y^{7.53}$). Estimates of these variables in the active $D_2R$ (PDB accession code: 6VMS) and $\beta_2AR$ (PDB accession code: 4LDO) crystal structures are represented by blue and green dots respectively. Estimates of these variables in the inactive $D_2R$ (PDB accession code: 6CM4) and $\beta_2AR$ (PDB accession code: 2RH1) crystal structures are represented by cyan and pink stars respectively. Projections of other class A GPCRs representative structures of active and inactive states are represented as dots and stars, respectively, including active $D_1$ dopamine receptor (PDB accession code: 7JVP) and $\beta_2AR$ (PDB accession code: 3SN6) and inactive $D_3$ dopamine receptor (PDB accession code: 3PBL) and $D_4$ dopamine receptor (PDB accession code: 5WIV).

**Table S1.** Total simulation time of each system in the enhanced sampling calculations.

| Ligand | PDB structure | Conventional MD | Steered MD | String method with swarms of trajectories/iterations | Total simulation time (μs) |
|---|---|---|---|---|---|
| Bromocriptine (crystalized) | 6VMS (active) | 500 ns | | | 0.5 |
| Risperidone (crystalized) | 6CM4 (inactive) | 500 ns | | | 0.5 |
| Apo | 6VMS (active) | | 102 ns | 400 (2.24 μs) | 2.342 |
| N-methyltyramine | 6VMS (active) | | 102 ns | 400 (2.24 μs) | 2.342 |
| 3-Methoxytyramine | 6VMS (active) | | 102 ns | 800 (4.48 μs) | 4.582 |
| Serotonin | 6VMS (active) | | 102 ns | 500 (2.8 μs) | 2.902 |
| p-Tyramine | 6VMS (active) | | 102 ns | 800 (4.48 μs) | 4.582 |
| m-Tyramine | 6VMS (active) | | 102 ns | 500 (2.8 μs) | 2.902 |
| Adrenaline | 6VMS (active) | | 102 ns | 400 (2.24 μs) | 2.342 |
| N-methyldopamine | 6VMS (active) | | 102 ns | 400 (2.24 μs) | 2.342 |
| Dopamine | 6VMS (active) | | 102 ns | 500 (2.8 μs) | 2.902 |
| Noradrenaline | 6VMS (active) | | 102 ns | 400 (2.24 μs) | 2.342 |
| Apo | 4LDO (active) | | 102 ns | 400 (2.24 μs) | 2.342 |
| Adrenaline (crystalized) | 4LDO (active) | | 102 ns | 400 (2.24 μs) | 2.342 |
| Dopamine | 4LDO (active) | | 102 ns | 400 (2.24 μs) | 2.342 |

**Table S2.** Collective variables for steered and string simulations of $D_2R$, which were obtained using the Demystify toolkit.

| Residues | Importance |
|---|---|
| V55 (1.53) - L375 (6.37) | 1.0 |
| W100 (23.50) - C385 (6.47) | 0.83 |
| L86 (2.56) - L375 (6.37) | 0.76 |
| I105 (3.23) - N175 (ECL2) | 0.74 |
| L375 (6.37) - Y426 (7.53) | 0.71 |
| V87 (2.57) - C401 (ECL3) | 0.67 |
| V47 (1.45) - G415 (7.42) | 0.67 |
| V191 (5.40) - S409 (7.36) | 0.67 |
| H106 (3.24) - C401 (ECL3) | 0.67 |
| G173 (4.63) - T412 (7.39) | 0.64 |
| V378 (6.40) - V421 (7.48) | 0.64 |
| N180 (ECL2) -D400 (ECL3) | 0.62 |
| C182 (45.50) - F189 (5.38) | 0.62 |
| A188 (5.37) - N396 (6.58) | 0.61 |
| T39 (1.37) - I403 (ECL3) | 0.60 |
| L41 (1.39) - E181 (ECL2) | 0.60 |
| K101 (23.51) - I394 (6.56) | 0.60 |
| R119 (5.68) - T412 (7.39) | 0.57 |
| N176 (ECL2) - A410 (7.37) | 0.57 |
| I183 (45.51) - C399 (6.61) | 0.55 |
| N35 (1.33) - E95 (2.65) | 0.55 |
| L41 (1.39) - L414 (7.41) | 0.53 |
| L44 (1.42) - N396 (6.58) | 0.53 |

| | |
|---|---|
| I73 (2.43) - Y416 (7.43) | 0.53 |
| M88 (2.58) - N180 (ECL2) | 0.52 |
| Y93 (2.62) - Q179 (ECL2) | 0.52 |
| V190 (5.39) - W413 (7.40) | 0.52 |
| I122 (3.40) - Y416 (7.43) | 0.52 |
| N124 (3.42) - V190 (5.39) | 0.52 |
| C182 (45.50) - H398 (6.60) | 0.52 |
| A188 (5.37) - Y416 (7.43) | 0.52 |
| F382 (6.44) - N422 (7.49) | 0.51 |
| R132 (3.50) - E368 (6.30) | |

**Table S3.** Local functional microswitches used to characterize the free energy landscapes

| Name | Measurement |
|---|---|
| RMSD of CWxP | RMSD of $C^{6.47}$, $W^{6.48}$, and $P^{6.50}$ heavy atoms to the active structure 6VMS |
| RMSD of PIF | RMSD of $I^{3.40}$ and $F^{6.44}$ heavy atoms to the active structure 6VMS |
| RMSD of NPxxY | RMSD of $N^{7.49}$, $P^{7.50}$ and $Y^{7.53}$ heavy atoms to the active structure 6VMS |
| TM3-TM6 distance | $R^{3.50}$ – $E^{6.30}$ Cα distance |
| TM5 bulge | $S^{5.46}$ – $V^{2.53}$ Cα distance |

**Table S4.** Relative experimental and calculated binding free energies (kcal/mol) for the $D_2R$ ligands.

| | Calculated active $\Delta\Delta G^a$ | Calculated inactive $\Delta\Delta G^a$ | Calculated $\Delta\Delta\Delta G^a$ | $pK_i^b$ | $E_{max}$ (%)$^b$ |
|---|---|---|---|---|---|
| Dopamine | - | - | - | 6.47 ± 0.50 | 100 ± 7 |
| Serotonin | - | - | - | 5.56 ± 0.43 | 7 ± 4 |
| p-Tyramine | 2.65 ± 0.32 | 0.39 ± 0.31 | 2.24 ± 0.20 | 4.11 ± 0.48 | 33 ± 5 |
| Adrenaline | 1.35 ± 0.66 | -0.02 ± 0.50 | 1.36 ± 0.67 | 5.15 ± 0.47 | 84 ± 5 |
| Noradrenaline | 1.53 ± 0.26 | 0.73 ± 0.42 | 0.87 ± 0.12 | 5.02 ± 0.47 | 98 ± 7 |
| m-Tyramine | 0.30 ± 0.12 | -1.25 ± 0.17 | 1.56 ± 0.04 | 5.73 ± 0.49 | 63 ± 4 |
| Hydroxyphenethylamine | 4.92 ± 0.38 | 0.77 ± 0.23 | 4.18 ± 0.15 | 3.71 ± 0.49 | 29 ± 3 |
| 3-Methoxytyramine | 1.25 ± 0.32 | -2.93 ± 0.43 | 4.14 ± 0.28 | 4.58 ± 0.48 | 13 ± 2 |
| Phenylephrine | 1.72 ± 0.36 | -1.94 ± 0.24 | 3.62 ± 0.18 | 4.82 ± 0.48 | 28 ± 6 |
| Norphenylephrine | 2.16 ± 0.22 | 0.15 ± 0.22 | 2.05 ± 0.06 | 4.77 ± 0.49 | 40 ± 6 |
| Methyldopamine | -0.56 ± 0.32 | -1.87 ± 0.22 | 1.13 ± 0.14 | 7.22 ± 0.49 | 96 ± 6 |
| N-Methyl-p-Tyramine | 2.15 ± 0.50 | -0.99 ± 0.37 | 3.12 ± 0.37 | 5.65 ± 0.52 | 6 ± 2 |
| Octopamine | 3.40 ± 0.46 | 0.68 ± 0.42 | 2.82 ± 0.35 | 3.46 ± 0.60 | 9 ± 8 |
| Phenethylamine | 3.57 ± 0.35 | 0.01 ± 0.11 | 3.55 ± 0.12 | 4.73 ± 0.52 | 28 ± 5 |
| N-Methylphenethylamine | 2.51 ± 0.44 | -1.33 ± 0.32 | 3.86 ± 0.24 | 4.77 ± 0.49 | 30 ± 4 |
| Synephrine | 2.95 ± 0.72 | -0.28 ± 0.41 | 3.30 ± 0.35 | 4.06 ± 0.51 | 3 ± 2 |

[a] Relative binding free energies compared to dopamine. Error bars represent the standard error of the mean (SEM) of five independent calculations

[b] Experimental binding affinities ($pK_i$) and $E_{max}$ values. Values represent the mean ± SEM of three individual experiments consisting of at least duplicates.
